# A novel factor essential for unconventional secretion of chitinase Cts1

**DOI:** 10.1101/2020.02.07.938613

**Authors:** Michèle Reindl, Janpeter Stock, Kai P. Hussnaetter, Aycin Genc, Andreas Brachmann, Kerstin Schipper

**Affiliations:** Institute for Microbiology, Heinrich Heine University Düsseldorf, Universitätsstraße 1, Düsseldorf, Germany; Bioeconomy Science Center (BioSC), Forschungszentrum Jülich, Jülich, Germany; Ludwig-Maximilians-Universität München, Faculty of Biology, Genetics, Planegg-Martinsried, Germany

**Author notes:** These authors contributed equally to this work (Michèle Reindl, Janpeter Stock).

**Keywords:** forward genetic screen, β-galactosidase, β-glucuronidase, unconventional secretion, *Ustilago maydis*, UV mutagenesis

## Abstract

Subcellular targeting of proteins is essential to orchestrate cytokinesis in eukaryotic cells. During cell division of *Ustilago maydis*, for example, chitinases must be specifically targeted to the fragmentation zone at the site of cell division to degrade remnant chitin and thus separate mother and daughter cells. Chitinase Cts1 is exported to this location via an unconventional secretion pathway putatively operating in a lock-type manner. The underlying mechanism is largely unexplored. Here, we applied a forward genetic screen based on UV mutagenesis to identify components essential for Cts1 export. The screen revealed a novel factor termed Jps1 lacking known protein domains. Deletion of the corresponding gene confirmed its essential role for Cts1 secretion. Localization studies demonstrated that Jps1 colocalizes with Cts1 in the fragmentation zone of dividing yeast cells. While loss of Jps1 leads to exclusion of Cts1 from the fragmentation zone and strongly reduced unconventional secretion, deletion of the chitinase does not disturb Jps1 localization. Yeast-two hybrid experiments suggest that the two proteins interact. In essence, we identified a novel component of unconventional secretion that functions in the fragmentation zone to enable export of Cts1. We hypothesize that Jps1 acts as an anchoring factor, supporting the proposed novel lock-type mechanism of unconventional secretion.

## 1. Introduction

Protein targeting is required to orchestrate essential cellular functions. Eukaryotic cells particularly rely on this process because of their compartmentalization and the necessity of equipping membrane-enclosed organelles with cognate protein subsets (Sommer and Schleiff, 2014). Protein targeting is mediated mostly by signal sequences. This is exemplified by the N-terminal signal peptide for entry of the endoplasmic reticulum (ER). The endomembrane system was thought to be the only export route for long time. However, recent years challenged this view by the finding that many proteins lacking a signal peptide are secreted by other mechanisms. The term unconventional secretion collectively describes protein export mechanisms that circumvent signal peptide-mediated passage through the canonical endoplasmic reticulum - Golgi pathway (Malhotra, 2013; Rabouille, 2017). Unconventional secretion has been discovered in lower eukaryotes like the fungal model *Saccharomyces cerevisiae* or the amoeba *Dictyostelium discoideum*, but also plays important roles in higher eukaryotes. It is even involved in human disease like in infections with the human immunodeficiency (HIV) or Epstein Barr viruses (Rayne et al., 2010; Debaisieux et al., 2012; Nowag and Münz, 2015). Research revealed that unconventional export mechanisms can be vesicular or non-vesicular (Rabouille et al., 2012), however, molecular details on the different pathways are scarce. The best described examples are self-sustained translocation of fibroblast growth factor 2 (FGF2) in human cells and the secretion of acyl-binding protein Acb1 via specialized compartments of unconventional secretion (CUPS) in *S. cerevisiae* (Malhotra, 2013; Steringer and Nickel, 2018).

Recently, a novel mechanism of unconventional secretion has been described for chitinase Cts1 in the model microorganism *Ustilago maydis* (Reindl et al., 2019). In its yeast form the fungus grows by budding. In these cells, Cts1 acts in concert with a second chitinase, Cts2, and mediates cell separation during cytokinesis. Elimination of both enzymes results in a cytokinesis defect and the formation of cell aggregates (Langner et al., 2015). Cts2 has a predicted N-terminal signal peptide and is thus thought to be secreted via the conventional secretion route, pointing towards an intricate interplay between both pathways.

In line with its cellular function, Cts1 translocates into the fragmentation zone of budding yeast cells (Langner et al., 2015; Aschenbroich et al., 2019). This unique small compartment arises between mother and daughter cell after consecutive formation of two septa at the cell boundary (Reindl et al., 2019). Recently we demonstrated that Cts1 release depends on cytokinesis and that the fragmentation zone is its most likely site of secretion, suggesting a lock-type mechanism (Aschenbroich et al., 2019). The septation proteins Don1 and Don3 are essential for Cts1 release (Aschenbroich et al., 2019). Don1 is a guanosine triphosphate exchange factor (GEF) that is delivered into the fragmentation zone by motile early endosomes, and Don3 is a germinal centre kinase (Weinzierl et al., 2002; Böhmer et al., 2009). Both proteins are required for secondary septum formation and their absence results in a cytokinesis defect similar to the one observed for the *cts1*/*cts2* deletion strain (Langner et al., 2015). To obtain further insights into subcellular targeting and unconventional secretion of Cts1, we here developed and applied a UV mutagenesis screen to identify components of the unconventional secretion pathway.

## 2. Materials and Methods

### Molecular biology methods

All plasmids (pUMa vectors, see below and Table 1) generated in this study were obtained using standard molecular biology methods established for *U. maydis* including Golden Gate cloning (Brachmann et al., 2004; Kämper, 2004; Terfrüchte et al., 2014; Bösch et al., 2016). Oligonucleotides applied for sequencing and cloning are listed in Table 2. Genomic DNA of strain UM521 (Kämper et al., 2006) was used as template for PCR reactions. All plasmids were verified by restriction analysis and sequencing. Detailed cloning strategies and vector maps will be provided upon request.

**Table 1.**
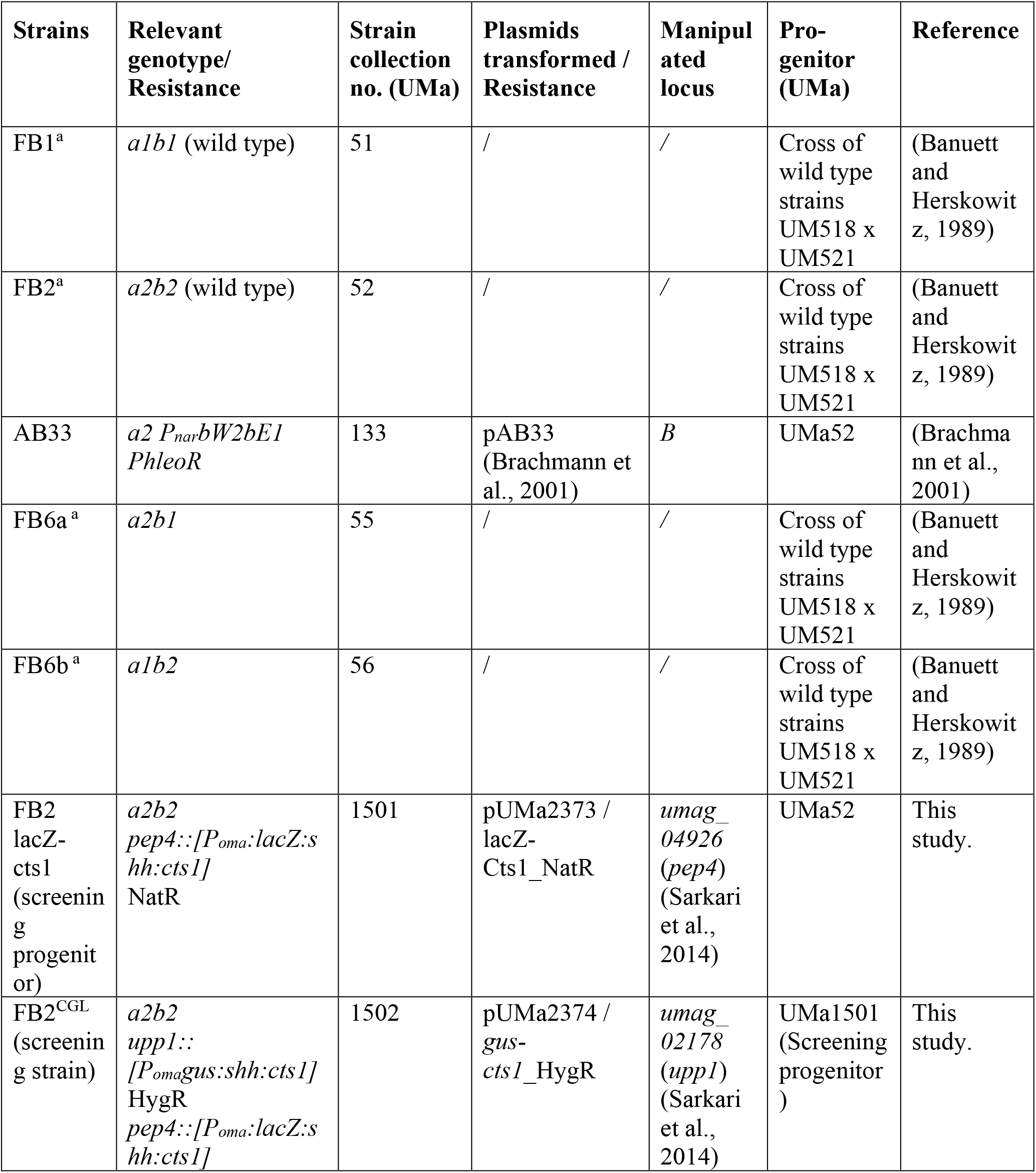
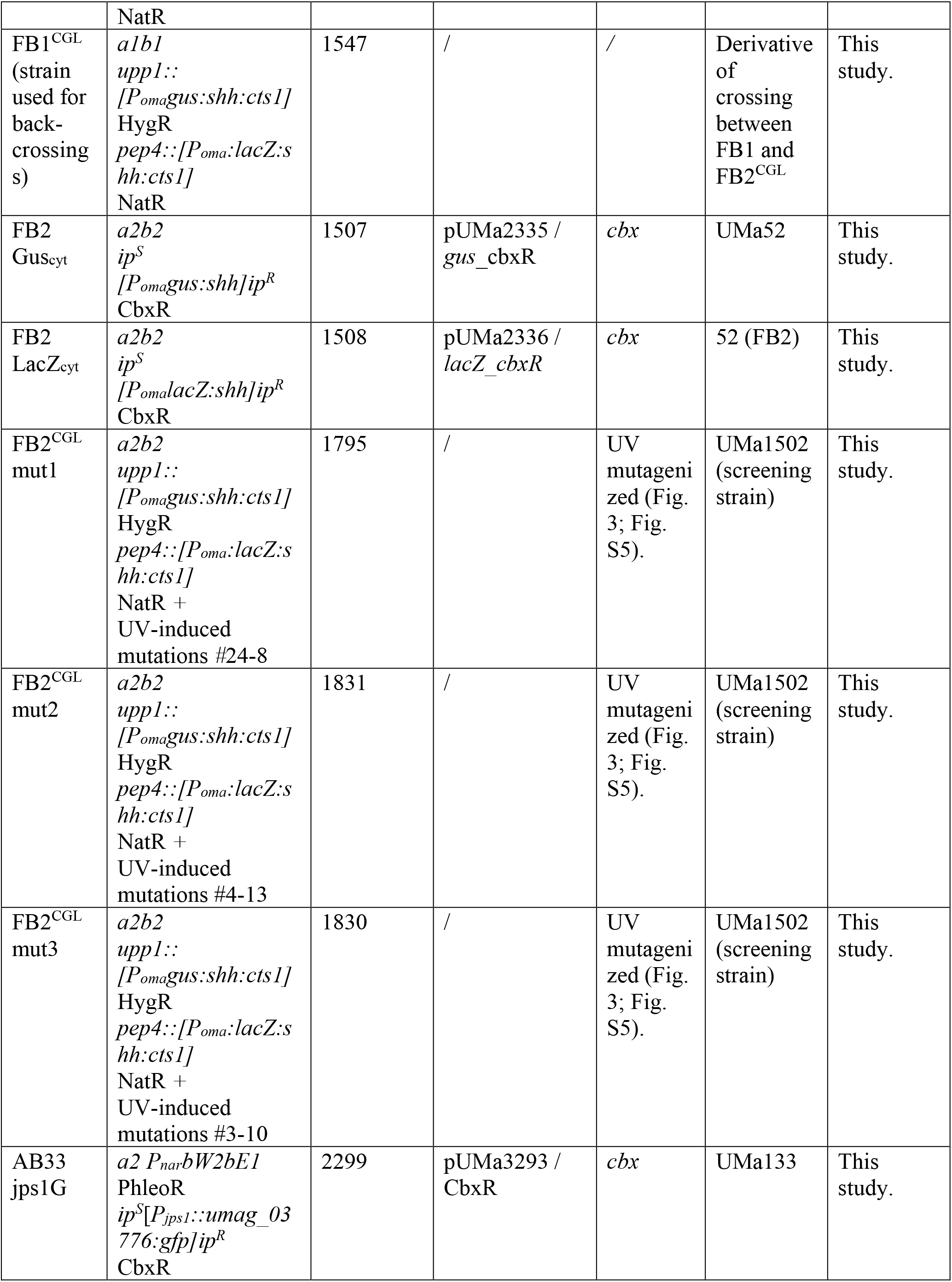
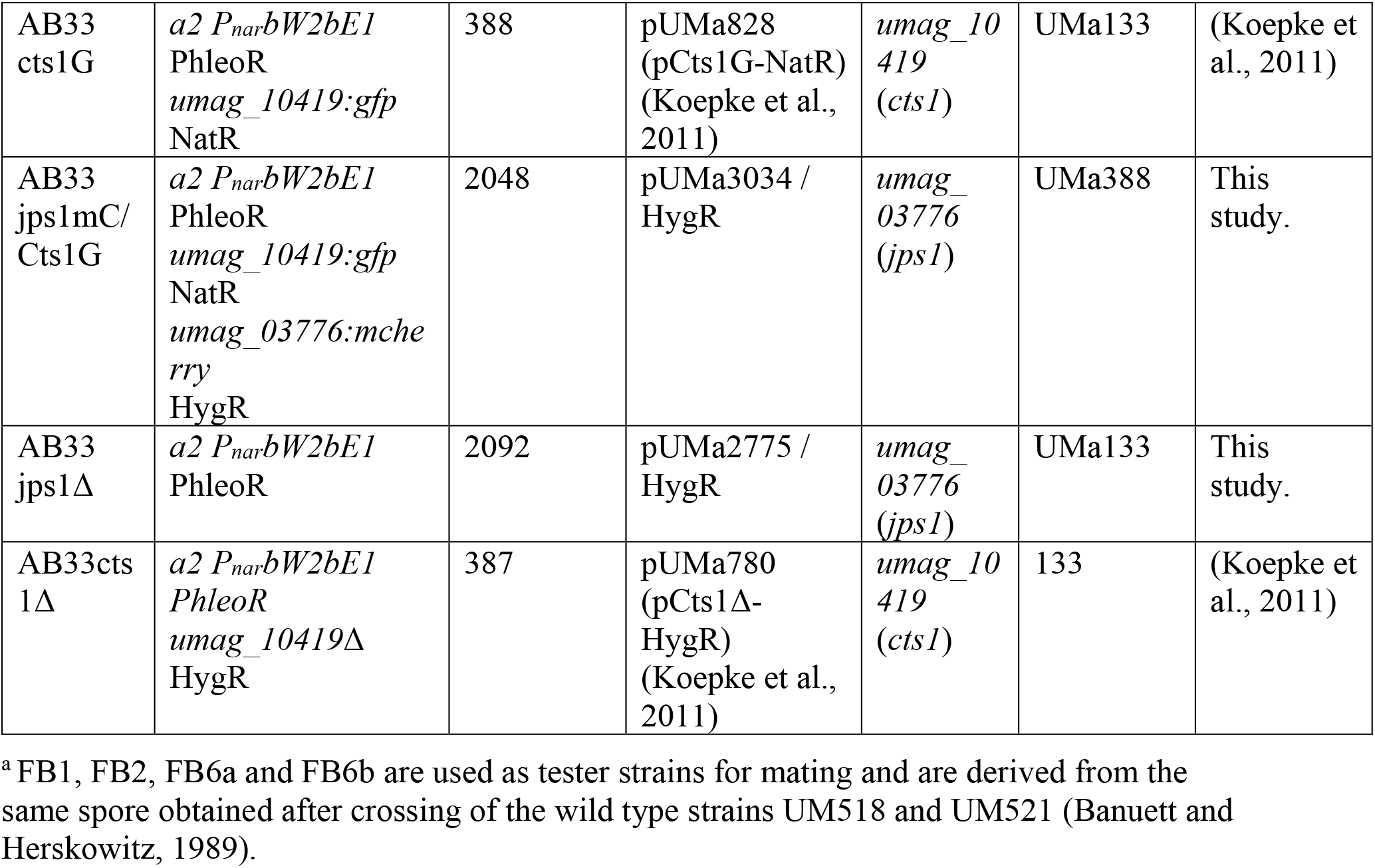
*U. maydis* strains used in this study.

**Table 2.**
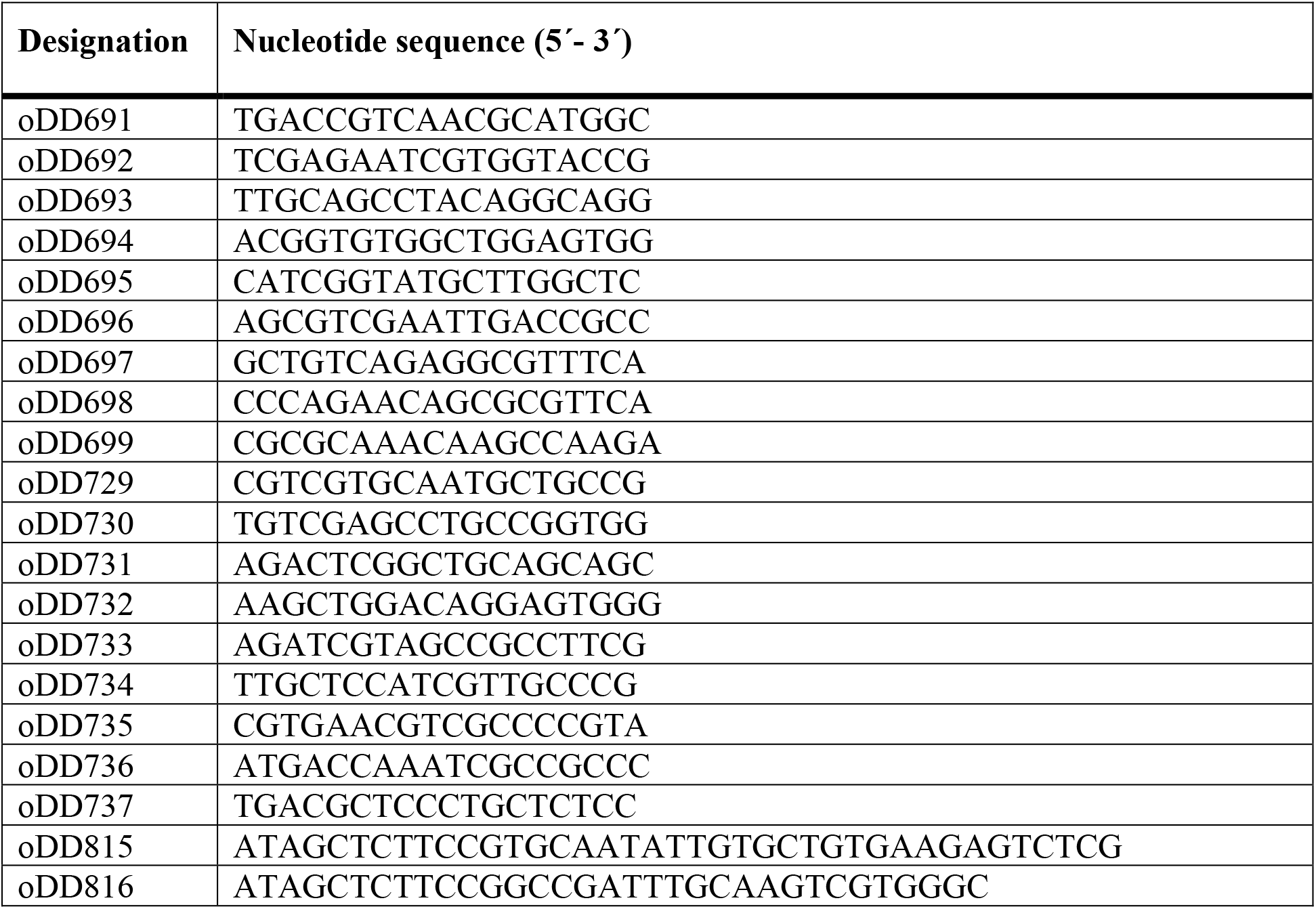
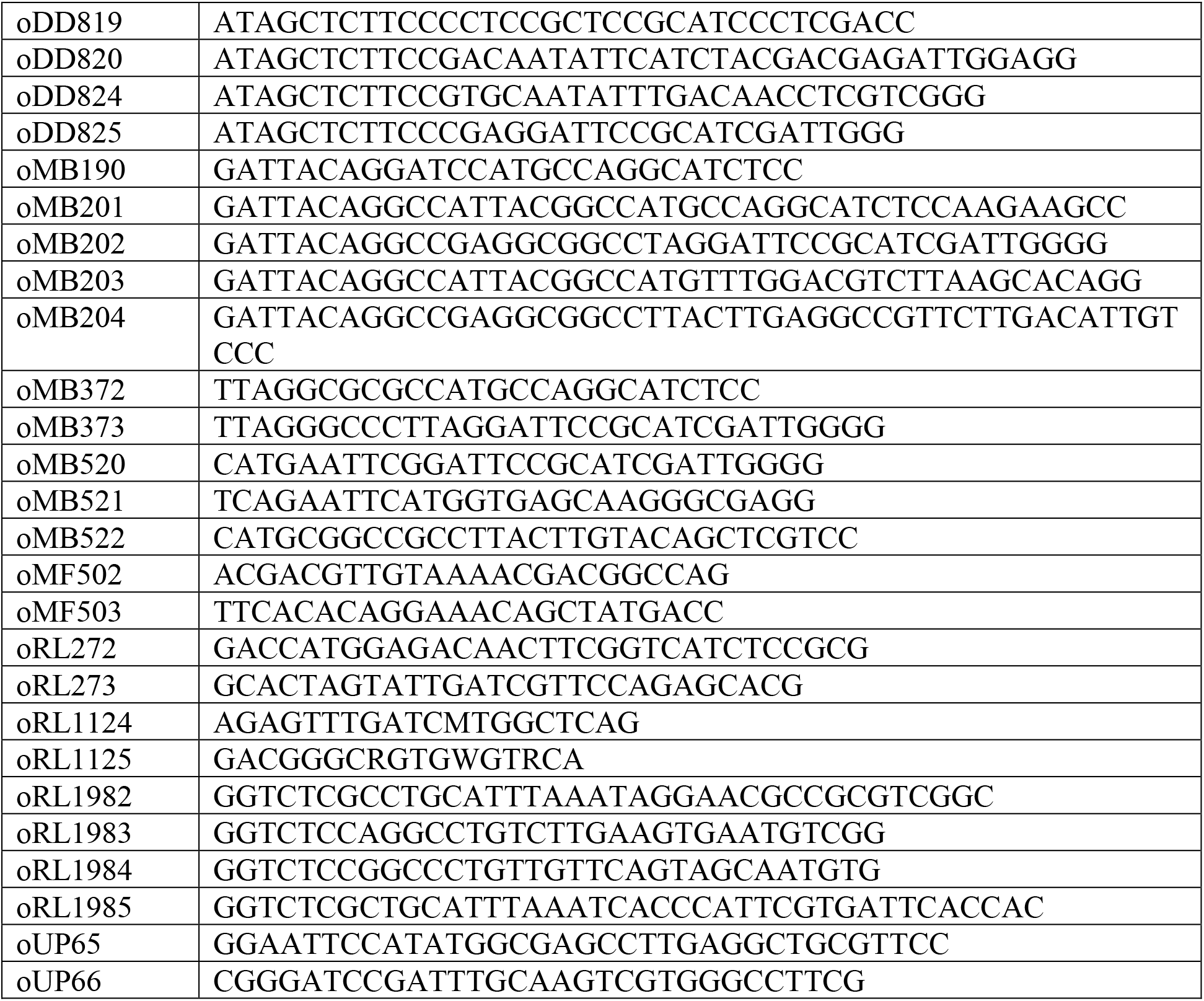
DNA oligonucleotides used in this study.

Plasmids for stable transformation of *U. maydis*: pUMa2373 (pDest-pep4D_Poma-LacZ-SHH-Cts1_NatR) was obtained in a Golden Gate cloning reaction using two flanking regions obtained by PCR using oligonucleotide combinations oRL1982 × oRL1983 (upstream flank *pep4* locus) and oRL1984 × oRL1985 (downstream flank *pep4* locus), the destination vector pUMa1476 (Terfrüchte et al., 2014) and pUMa2372. Storage vector pUMa2372 contained a P*oma* controlled non-optimized version of the *lacZ* gene (beta-D-galactosidase, accession NP_414878.1) from *Escherichia coli* Rosetta 2 in translational fusion to the *cts1* gene (*umag_10419*) via an SHH linker (Sarkari et al., 2014) and a nourseothricin-resistance marker cassette (NatR) flanked by BsaI sites. To generate pUMa2335 (pRabX1-Poma_Gus-SHH_cbx), pUMa2113 was hydrolyzed with NcoI and NotI and a Gus-SHH encoding fragment was inserted, replacing the previous insert coding for Gus-SHH-Cts1. Similary, in pUMa2336 (pRabX1-Poma_LacZ-SHH_cbx) the insert in pUMa2113 was replaced via NcoI/NotI restriction/ligation by a fusion gene encoding LacZ-SHH. Both plasmids were integrated in the *ip* locus under the control of the strong, constitutively active *oma* promoter. pUMa2605 was obtained by hydrolysis of p*cts2*∆_hyg (Langner et al., 2015) with SfiI and replacing the HygR cassette with the G418-resistance cassette (G418R) from pUMa1057 (pMF1g) (Baumann et al., 2012). For assembly of pUMa3012 (pRabX1-Poma_Gus-SHH-Jps1) the *jps1* gene was amplified by PCR using oMB372 × oMB373 yielding a 1844 bp product flanked by AscI and ApaI restriction sites. After hydrolysis with these enzymes the *jps1* gene replaced the *cts1* gene in pUMa2113 (Sarkari et al., 2014). For assembly of pUMa3034 flanking regions were amplified with oDD824 × oDD825 (upstream flank *umag_03776* gene) and oDD819 × oDD820 (3’region of *umag_03776*) and used for SapI mediated Golden Gate cloning including destination vector pUMa2074 and the storage vector pUMa3035. pUMa3035 contained the mCherry gene for translational fusions as well as a HygR cassette. pUMa3111 (pDest-jps1D_G418R) was generated by replacing the HygR cassette in pUMa2775 by a G418R cassette from pUMa1057 using flanking SfiI sites. The progenitor vector pUMa2775 (pDest-jps1D_HygR) was synthetized by SapI mediated Golden Gate cloning with flanking regions obtained by PCR with oDD815 × oDD816 (upstream flank for *umag_03776*) and oDD819 × oDD820 (downstream flank for *umag_03776*), destination vector pUMa2074 and storage vector pUMa2242 harboring a HygR cassette (Aschenbroich et al., 2019). For generation of pUMa3293 (pRabX1-Pjps1_Jps1_eGfp_CbxR) a 1952 bp PCR product obtained with oUP65 × oUP66 (*umag_03776* promoter region) was hydrolyzed with NdeI and BamHI and inserted into a pRabX1 derivative (pUMa3095) upstream of a *jps1*:*gfp* fusion gene (Stock et al., 2012). pUMa3095 was assembled in a three-fragment ligation of a 1849 bp PCR product of oMB190 × oMB120 (*umag_03776* gene) hydrolyzed with BamHI and EcoRI, a 741 bp PCR product of oMB521 × oMB522 (*gfp* gene) hydrolyzed with EcoRI and NotI and a 6018 bp fragment of vector pUMa2113 (Sarkari et al., 2014) hydrolyzed with BamHI and NotI.

Plasmids for two-hybrid analyses in *S. cerevisae*: All Yeast-two hybrid plasmids were generated on the basis of the Matchmaker III System (Clontech Laboratories Inc., Mountain View, CA, USA). pGAD and pGBKT7 were modified to contain a SfiI site with specific overhangs to exchange the gene of interest. For generation of pUMa2927 (pGAD_Jps1) and pUMa2929 (pGBK_Jps1) a 1869 bp PCR product was obtained with oMB201 and oMB202 (amplifying the *jps1* gene). The PCR product was hydrolyzed with SfiI and inserted into pGAD and pGBK backbones. For generation of pUMa2928 (pGAD_Cts1) and pUMa2930 (pGBK_Cts1), a 1549 bp PCR product was obtained with oMB203 and oMB204 (amplifying the *cts1* gene). The product was hydrolyzed with SfiI and inserted into pGAD and pGBK backbones.

### Strains and cultivation conditions

*U. maydis* strains used in this study were obtained by homologous recombination yielding genetically stable strains (Table 1). For genome insertion at the *ip* locus, integrative plasmids were used (Stock et al., 2012). These plasmids contain an *ip*^R^ allele that mediates carboxin resistance (Keon et al., 1991). Integrative plasmids were linearized within the *ip*^R^ and subsequently used to transform *U. maydis* protoplasts. Mutants harboring a single copy of the plasmid were obtained via homologous recombination (Brachmann et al., 2004; Kämper, 2004). Gene deletion and translational fusions *in locus* were performed with plasmids obtained by either the SfiI- or Golden Gate cloning strategy using plasmids deposited in the Institutes plasmid collection (Brachmann et al., 2004; Terfrüchte et al., 2014) http://www.mikrobiologie.hhu.de/ustilago-community.html). Gene insertion at the *pep4* (*umag_04926*) and *upp1* (*umag_02178*) locus resulted in the deletion of the respective protease-encoding genes and hence, as a positive side-effect for the screen diminished proteolytic activity of the strains (Sarkari et al., 2014). The corresponding plasmids were obtained by Golden Gate cloning (Terfrüchte et al., 2014). All strains generated were verified by Southern blot analysis using digoxygenin labeled probes (Roche). For *ip* insertions, the probe was obtained with the primer combination oMF502/oMF503 and the template pUMa260 (Loubradou et al., 2001). For insertions at the *pep4* or *upp1* locus (Sarkari et al., 2014) and other in-locus manipulations, the two flanking regions (upstream and downstream flanks) were amplified as probes.

*U. maydis* strains were grown at 28 °C in complete medium (CM) supplemented with 1% (w/v) glucose (CM-glc) (Holliday, 1974) or YepsLight (modified from (Tsukuda et al., 1988)). Solid media were supplemented with 2% (w/v) agar agar. CM-glc plates containing 1% (w/v) charcoal (Sigma C-9157) were used for mating assays (Hartmann et al., 1996).

*Saccharomyces cerevisiae* strain AH109 (Clontech Laboratories Inc., Mountain View, CA, USA) was employed for yeast two-hybrid assays. Gene sequences without predicted introns were inserted into the vectors pGAD24 and pGBKT7 generating translational fusions to the Gal4 activation domain (AD) and DNA-binding domain (BD), respectively.

### Generation of a compatible strain by genetic crossings

To enable genetic back-crosses with FB2^CGL^ the reporters were also introduced into the compatible strain FB1 by genetic crosses. Therefore, wild type strain FB1 was crossed with screening strain FB2^CGL^ (mating type *a2b2*) using plant infections (see below) to obtain meiotic progeny. These were tested for their ability to mate with FB2. Compatible mating was screened on CM-glc plates containing 1% (w/v) charcoal (Fig. S1) on which strains harboring different alleles of both mating type loci form fuzzy colonies while strains which do not mate grow in smooth colonies. First, progeny was screened for the presence of the two artificial reporters Gus-Cts1 and LacZ-Cts1. Positive candidates were then tested in mating experiments with tester strains for induction of the fuzzy phenotype in FB2 crosses.

### Mixing experiments to distinguish intra- and extracellular reporter activities

To distinguish intra- and extracellular LacZ activity defined cell amounts were mixed on CM-glc/X-Gal plates containing 1% (w/v) glucose and X-Gal (5-bromo-4-chloro-3-indolyl-β-D-galactopyranoside; 20 mg/ml in DMSO, f.c. 60 mg/L in CM-glc). Screening strain FB2^CGL^ and the AB33LacZ_cyt_ control were grown in liquid CM-glc medium until logarithmic phase. Cells were harvested, washed in PBS (1x, pH 7.2) and adjusted to an OD_600_ of 1.0. A 10^−4^ serial dilution was prepared in PBS. The diluted suspensions of both strains were mixed in defined ratios and plated in a total volume of 150 μl on CM-glc/X-Gal plates. After incubation for 4 days at 28 °C protected from light, growth and conversion of substrate was photographed. For illumination a Ledgo CN-B150 LED On-Camera Light was used.

### Gus/LacZ activity plate and membrane assays

Gus and LacZ activity were tested by indicator plate assays using CM plates containing 1% (w/v) glucose (CM-glc) and the respective chromogenic substrate X-Gluc (5-bromo-4-chloro-3-indolyl-beta-d-glucuronic acid; 0.5 mg/ml in DMSO) or X-Gal (20 mg/ml in DMSO), respectively. Tested strains were grown in CM-glc for 16 hours. After adjusting the cultures to an OD_600_ of 1.0 in sterile PBS, 10 μl suspension were spotted on CM-glc plates and incubated at 28 °C for 2 days. Intracellular Gus and LacZ activity was visualized by placing a nitrocellulose membrane (Amersham ^TM^ Protran TM 0.45 μM NC, GE Healthcare Life Sciences) on top for 24 h at 28 °C. The membrane was then removed and treated with liquid nitrogen for 3 min for cell lysis. Subsequently, it was soaked in X-Gluc buffer (25 μg X-Gluc/ml, solved in DMSO, 5 mM sodium phosphate buffer pH 7.0, 14 μM β-mercaptoethanol, 0.4 mM EDTA, 0.0021% (v/v) lauroyl-sarcosin, 0.002% (v/v) Triton X-100, 0.1 mg/ml (w/v) BSA) or X-Gal Buffer (1 mg X-Gal/ml, solved in DMSO, 15 mM Sodium phosphate buffer pH 7, 5 mM KCl, 0.5 mM MgSO4, 34 mM β-mercaptoethanol) for Gus and LacZ activity, respectively, and incubated at 37 °C for 18 h.

### UV mutagenesis

For UV mutagenesis a 20 ml YepsLight pre-culture of screening strain FB2^CGL^ was inoculated from a fresh plate and incubated overnight (200 rpm, 28 °C). In the morning, the culture was diluted to an OD_600_ of 0.1 in 20 ml (200 rpm, 28 °C). The culture was incubated until it reached an OD_600_ of 0.5. Subsequently it was diluted stepwise to an OD_600_ of 0.00125 (1: 400) in 20 ml YepsLight. 150 μl of the 1:400 dilutions were spread evenly onto CM-glc screening plates containing 10 μg/ml X-Gal. The dried plates were exposed to UV irradiation (30 mJ/cm^2^) using a Stratalinker device (Stratagene). Plate lids were removed during exposition. Subsequently, plates were incubated for 2 to 3 days at 28 °C until single colonies were grown.

### Screening for diminished reporter secretion

Clones that showed reduced or absent LacZ activity (i.e., colorless appearing colonies) after UV mutagenesis on CM-glc screening plates containing X-Gal (see above) were patched on plates containing X-Gluc to additionally assay for extracellular Gus activity. Plates were prepared by spreading 100 μl X-Gluc solution (100 mg/ml stock in DMSO) on CM-glc plates. Plates were incubated for 2 to 3 days on 28 °C. Colorless colonies were patched again on X-Gluc and X-Gal plates simultaneously, this time streaking out larger areas of about 0.5 × 0.5 cm. During the procedure, control strains producing intracellular LacZ or Gus (FB2 LacZ_cyt_ and FB2 Gus_cyt_, respectively), the non-mutagenized screening strain (FB2^CGL^) and the precursor strain lacking any reporters (FB2) were handled in parallel to verify the results (Table 1).

### Generation of cell extracts and supernatants

Strains were inoculated in 20 ml CM-glc and incubated at 28 °C overnight. Next morning, the culture was used to inoculate a new culture of 70 ml CM-glc with a starting OD_600_ of 0.05. To detect potential growth defects, growth of the culture was followed by determining the OD_600_ every hour for at least 8 to 10 hours. 2 ml aliquots of supernatants for Gus/LacZ assays and whole cells for Cts1 assays were harvested at OD_600_ of 0.3. Once the culture reached an OD_600_ of 0.7 50 ml were harvested (5 min, 3000 rpm, 4 °C). The supernatant was transferred to a new tube and stored on 4 °C. The cell pellet was used to prepare native cell extracts used for the Cts1, Gus and LacZ assay (modified from (Stock et al., 2016)). To this end the cell pellet was resuspended in 2 ml ice-cold native extraction buffer (1 mM phenylmethylsulfonylfluorid (PMSF), 2.5 mM benzamidine hydrochloride hydrate, 1 μM pepstatin; 100 μL Roche EDTA-free protease inhibitor cocktail 50×; dissolve in PBS pH 7.4). The suspension was then frozen in pre-chilled metal pots (25 ml, Retsch) using liquid nitrogen. Cells were ruptured at 4 °C using the Retsch mill (10 min, 30 Hz) and then the metal pots were thawed at 4 °C for 1 hour. The cell extracts were transferred to a reaction tube and centrifuge for 30 min (4 °C, 13000 rpm, benchtop centrifuge). Bradford assays were conducted to determine the protein concentrations in the samples (Bradford, 1976).

### Quantitative determination of Gus and LacZ activity

Importantly, quantitative Gus, LacZ and Cts1 assays were conducted from a single culture (see below for Cts1 assay). Quantitative Gus and LacZ liquid assays were based on the chromogenic and fluorescent substrates ONPG for the liquid LacZ activity assay (o-nitrophenyl-β-D-galactopyranoside) and MUG for the liquid Gus activity assay (4-methylumbelliferyl-ß-D-glucuronide trihydrate; BioWorld, 30350000-2 (714331), respectively (Stock et al., 2012; Stock et al., 2016). For both the ONPG and the MUG liquid assays (modified from (Stock et al., 2012; Stock et al., 2016)), activity was determined in native cell extracts and in the cell-free culture supernatant of candidate mutants in comparison to control strains (Koepke et al., 2011; Stock et al., 2012; Langner et al., 2015).

The Gus and LacZ assays with cell extracts and cell-free supernatants were conducted according to slightly modified published protocols (Miller, 1959; Stock et al., 2012). To this end, native cell extracts were adjusted to a total protein concentration of 100 μg/ml using PBS buffer. 10 μl of native cell extracts were then mixed in a black 96-well plate (96 Well, PS, F-Bottom, μCLEAR, black, CELLSTAR) with 90 μl of Gus- or Z-buffer and 100 μl of the respective substrate solution (Gus: 2 mM MUG, 1/50 vol. bovine serum albumin fraction V (BSA) in 1x Gus buffer; LacZ: 1 mg/ml ONPG in 2x Z-buffer). For supernatant measurements, 100 μl cell-free supernatants were mixed with 100 μl of the respective substrate solution. 2x Gus buffer (Stock et al., 2016) was used for the Gus assay and 2x Z-buffer (80 mM Na2HPO_4_, 120 mM NaH_2_PO_4_*H_2_O, 20 mM KCl, 2 mM MgSO_4_*7H_2_O; adjust to pH 7; add 100 mM β-mercaptoethanol freshly) was used for the LacZ assay. The assays were conducted in the Tecan device (Tecan Group Ltd., Männedorf, Switzerland) for 1 h at 37 °C with measurements every 10 min (excitation/emission wavelengths: 365/465 nm for Gus activity; OD_420_ for LacZ activity). A fixed gain of 150 was used for cell extract measurements and fixed values of 60 and 100 for Gus and Cts1 activity assays of culture supernatants, respectively.

For data evaluation the slope during linear activity increase of the kinetic measurements was determined. Values for the screening strain FB2^CGL^ were set to 100% to judge the activities in the mutants.

### Quantitative determination of Cts1 activity

The fluorescent substrate MUC was applied for the Cts1 liquid assay (4-methylumbelliferyl β-D-N,N′,N′′-triacetylchitotrioside hydrate; M5639 Sigma-Aldrich) (Koepke et al., 2011; Stock et al., 2012). Strain AB33 cts1Δ (UMa387) (Koepke et al., 2011) carrying a *cts1* deletion dealt as negative control for the Cts1 assays. For the MUC assay whole cells were subjected to the assay after washing to detect Cts1 activity at the cell surface. For intracellular activities, cell extracts were used (see above).

The Cts1 activity liquid assay with whole cells or 10 μg of native cell extracts was conducted according to published protocols with minor changes (Koepke et al., 2011; Stock et al., 2012). Once the culture had reached an OD_600_ of 0.3 a 2 ml sample was taken and cells were harvested by centrifugation (3 min, 8000 rpm, bench-top centrifuge). Cells were resuspended in 1 ml KHM buffer (110 mM potassium acetate, 20 mM HEPES, 2 mM MgCl_2_) (Koepke et al., 2011; Stock et al., 2012), the OD_600_ was documented and the suspension was subjected to the MUC assay. A MUC working solution was prepared form a stock solution (2 mg/ml MUC in DMSO) by diluting it 1:10 with KHM buffer (protect from light, store at 4 °C). Black 96-well plates (96 Well, PS, F-Bottom, μCLEAR, black, CELLSTAR) were used for the assay. 70 μl of working solution were mixed with 30 μl of the cell suspension in one well. Activities for each strain were determined in triplicates. The plates were sealed with parafilm and incubated in the dark for 1 h at 37 °C. The reaction was then stopped by adding 200 μl 1 M Na_2_CO_3_ and relative fluorescence units were determined in a plate reader at excitation and emission wave length of 360/450 nm, respectively, at 37 °C with a fixed gain of 100 (Tecan Reader).

### Plant infection and genetic back-crosses

Compatible *U. maydis* strains were subjected to genetic crosses on corn plants. Strain FB1^CGL^ was obtained by genetic crossing of FB1 (mating type *a1b1*) with FB2^CGL^ (mating type *a2b2*). Furthermore, mutagenized strains (derived from the FB2^CGL^ strain background; mating type *a2b2*) which showed strongly reduced Cts1 secretion assayed by all three reporters were subjected to back-crosses with the compatible wild type strain FB1^CGL^ (mating type *a1b1*). For infection of the host plant *Z. mays* (Early Golden Bantam), strains were grown to an OD_600_ of 0.8 in CM-glc, washed three times with H_2_O, and resuspended to an OD_600_ of 1 in H_2_O. For infection, compatible strains were then mixed in a 1:1 ratio. The cell suspension was injected into seven day-old maize seedlings. Virulence of strain crosses was quantified using established pathogenicity assays (Kämper et al., 2006). Therefore, 7 days post infection plants were scored for symptom formation according to the following categories: (1) no symptoms, (2) chlorosis, (3) anthocyanin accumulation, (4) small tumors (<1 mm), (5) medium tumors (>1 mm), and heavy tumors associated with bending of stem. For spore collection, mature tumor material was harvested two to three weeks after infection and dried at 37 °C for about 7 days.

### Spore germination and analysis of progeny

Spore germination and analysis was conducted according to published protocols (Eichhorn et al., 2006). To germinate spores for progeny analysis after genetic back-crossing of FB1 × FB2^CGL^ or FB1^CGL^ × FB2^CGL^mut1 dried tumor material was homogenized in a mortar, treated with 2 ml of a solution of 3.0% (w/v) copper sulfate for 15 min and washed twice with 1 ml sterile H_2_O. The spores were then resuspended in 500 μl sterile water. Prior to plating, the spore solutions were supplemented with ampicillin and tetracyclin to avoid bacterial contaminations (final concentrations: 600 μg/ml ampicillin; 150 μg/ml tetracyclin). Then, 200 μl of 1:1, 1:10 and 1:100 dilutions were spread on CM-glc plates, and incubated for 2 d at 28 °C. Resulting colonies were singled out again on CM-glc plates to guarantee that each colony results from one clone. To identify FB1^CGL^ after crossing of FB1 × FB2^CGL^ progeny was assayed on charcoal plates for their mating types (see below) and on X-Gal and X-Gluc plates for the presence of the two reporters Gus and LacZ. To analyze FB1^CGL^ and FB2^CGL^mut1 again indicator plates containing X-Gal and X-Gluc were used to pre-sort the cells into secretion competent and deficient clones. Candidates were further tested using liquid assays for all three reporters.

### Mating assay

For mating assays, cells were grown in CM medium to an OD_600_ of 1.0 and washed once with sterile H_2_O. Washed cells were adjusted to an OD_600_ of 3 in sterile H_2_O. Indicated strains were pre-mixed in equal amounts and then co-spotted on CM-glc plates containing 1% (w/v) charcoal. Plates were incubated at 28 °C for 24 h. Tester strains were used as controls (Table 2).

### Genome sequencing and assembly

Genomic DNA extraction was performed according to published protocols (Bösch et al., 2016), two separate preparations were combined, and the DNA concentration was adjusted to 100 ng/μl using TE-RNase in approximately 500 μl final volume. gDNA quality was verified by PCR reactions using primers specific for the bacterial 16sRNA gene (oRL1124 × oRL1125) and the intrinsic gene *uml2* (*umag_01422*; oRL272 × oRL273). For genome sequencing, DNA libraries were generated using the Nextera XT Kit (Illumina) according to manufacturer's instructions. Sequencing (v3 chemistry) was performed with a MiSeq sequencer (Illumina) at the Genomics Service Unit (LMU Biocenter). Obtained reads were quality trimmed and filtered with trimmomatic version 0.30 and fasta toolkit version 0.13.2. Sequence assemblies with passed single and paired reads were performed using CLC Genomics Workbench 8.0 (QIAGEN). The genome sequence of UM521 (*Ustilago maydis* 521; http://www.ncbi.nlm.nih.gov/nuccore/AACP00000000.2; accessed on 2020/04/01) was used as template for read assembly. Genomic DNA of screening strain FB2^CGL^ was sequenced as reference.

### Identification of genomic mutations

For identification of the underlying mutation in progeny of FB1^CGL^ and FB2^CGL^mut1, PCR products were generated from gDNA of progeny clones #3 and #5 which showed no Cts1 activity and no blue halos on X-Gal and X-Gluc indicator plates. To this end, the following primer combinations were used to amplify the indicated genes (Table 2): *umag_00493*: oDD691 × oDD729; *umag_06269*: oDD692 × oDD730; *umag_02631*: oDD693 × oDD731; *umag_03776*: oDD694 × oDD732; *umag_04298*: oDD695 × oDD733; *umag_05386*: oDD696 × oDD734; *umag_04385*: oDD697 × oDD735; *umag_04494*: oDD698 × oDD736 and *umag_11876*: oDD699 × oDD737. gDNA was obtained like described (Bösch et al., 2016). Additional sequencing was conducted for *umag_06269* and *umag_03776* using progeny #9, #26, #31 and #39 (not shown). PCR products were sequenced and analyzed for the presence or absence of the mutations identified in genome sequence alignments of FB2^CGL^ and FB2^CGL^mut1.

### Protein precipitation from culture supernatants

Secreted proteins were enriched from supernatant samples using trichloric acid (TCA) precipitation. Therefore, culture supernatants were supplemented with 10% (w/v) TCA and incubated overnight on 4 °C. After washing twice in −20 °C acetone the protein pellets were resuspended in minimal amounts of 3x Laemmli-Buffer (Laemmli, 1970) and the pH was eventually neutralized with 1 M NaOH. For SDS-Page analysis the samples were first boiled for 10 min and then centrifuged (22,000 × g, 5 min, room temperature).

### SDS-Page and Western blot analysis

Boiled protein samples were separated by SDS-Page using 10% (w/v) acrylamide gels. Subsequently, proteins were blotted to methanol-activated PVDF membranes. The analyzed proteins contained an SHH-tag (Sarkari et al., 2014) and were detected using anti-HA (Sigma-Aldrich, USA) antibodies and anti-mouse IgG-HRP (Promega, USA) conjugates as primary and secondary antibodies, respectively. HRP activity was detected using AceGlow Western blotting detection reagent (PeqLab, Germany) and a LAS4000 chemiluminescence imager (GE LifeScience, Germany).

### Yeast-two hybrid assays

Yeast two-hybrid analysis was carried out using the Clontech MatchMaker III system as described before (Pohlmann et al., 2015). Transformation with plasmids and cultivation were performed using standard techniques (Clontech manual). In addition to negative and positive controls included in the MatchMaker III system, also examples for weakly interacting proteins were included (Pohlmann et al., 2015). Auto-activation was excluded for all tested proteins using controls with control plasmids.

### Microscopy, image processing and staining procedures

Microscopic analysis was performed with a wide-field microscope from Visitron Systems (Munich, Germany), Zeiss (Oberkochen, Germany) Axio Imager M1 equipped with a Spot Pursuit CCD camera (Diagnostic Instruments, Sterling Heights, MI) and objective lenses Plan Neofluar (40x, NA 1.3) and Plan Neofluar (63x, NA 1.25). Fluorescence proteins were excited with an HXP metal halide lamp (LEj, Jena, Germany) in combination with filter sets for Gfp (ET470/40BP, ET495LP, ET525/50BP), mCherry (ET560/40BP, ET585LP, ET630/75BP, Chroma, Bellow Falls, VT), and DAPI (HC387/11BP, BS409LP, HC 447/60BP; AHF Analysentechnik, Tübingen, Germany). The system was operated with the software MetaMorph (Molecular Devices, version 7, Sunnyvale, CA). Image processing including adjustments of brightness and contrast was also conducted with this software. To visualize fungal cell walls and septa 1 ml of cell culture was stained with calcofluor white (1 μg/ml) before microscopy.

## 3. Results

### A genetic screen for mutants impaired in Cts1 secretion identifies a novel component

To identify mutants impaired in Cts1 secretion, a forward genetic screen based on UV mutagenesis was established (Fig. 1, Fig. S2; for experimental details see Materials and Methods section). To this end, a screening strain derived from the haploid FB2 wild type strain (mating type *a2b2*) was developed. This allows for co-infections of the host plant maize with compatible strains of differing *a* and *b* alleles like FB1 (mating type *a1b1*) to obtain meiotic progeny (Banuett and Herskowitz, 1989).

**Fig. 1.**
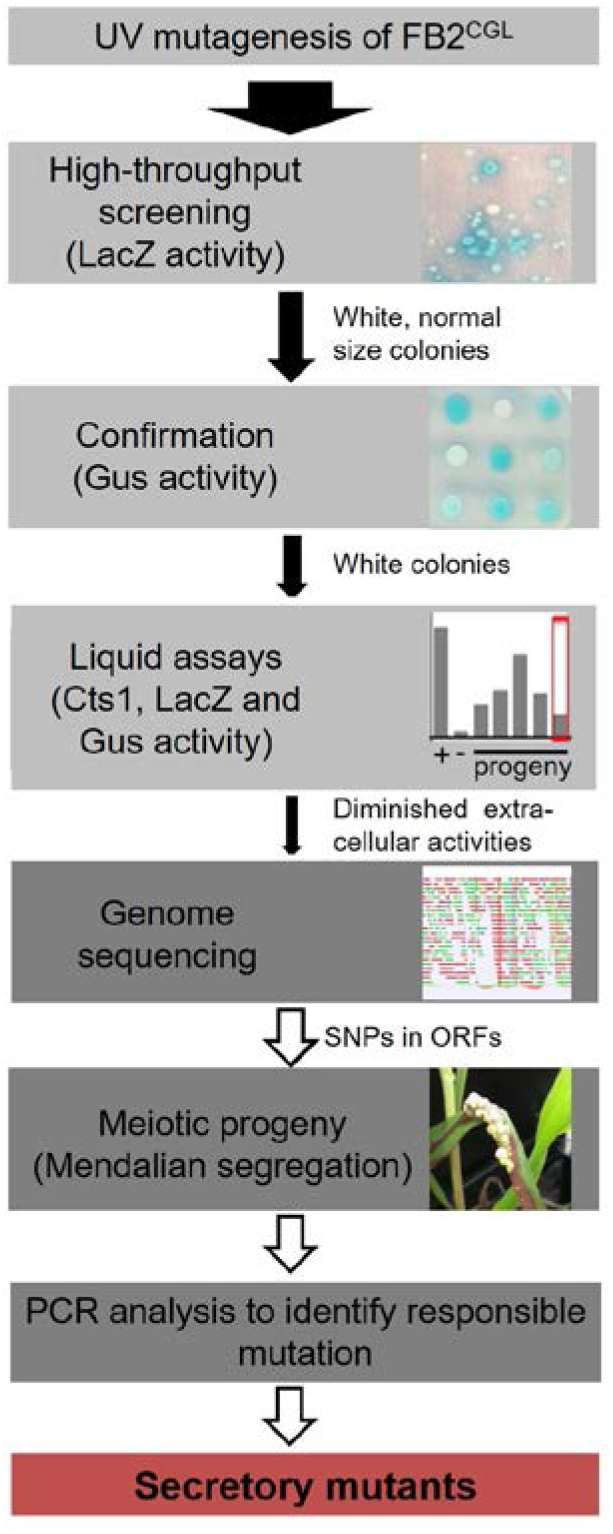
Rationale of the forward genetic screen. The three reporters LacZ-Cts1, Gus-Cts1 and endogenous Cts1 were used to identify mutants with diminished Cts1 secretion. After UV mutagenesis, a high-throughput screen for absence of LacZ activity of LacZ-Cts1 was conducted on X-Gal containing plates. Next, colonies were patched on X-Gluc containing plates to verify the result with the Gus-Cts1 reporter. Remaining candidates were assayed in quantitative liquid assays for extra- and intracellular LacZ, Gus, and Cts1 activity. Candidates with diminished extracellular activity of all three reporters but unimpaired intracellular activity were collected. After genome sequencing, the mutation responsible for diminished secretion was identified by PCR analysis of different meiotic progeny showing a similar secretion phenotype on loci containing SNPs as identified in the genome comparison with the progenitor strain. For details see Fig. S2.

To minimize false positive screening hits, three different reporters for Cts1 secretion were employed: the bacterial reporter enzymes β-glucuronidase (Gus; published in (Stock et al., 2012; Stock et al., 2016)) and β-galactosidase (LacZ; newly established for *U. maydis*) as fusion proteins with Cts1 (Gus-Cts1, LacZ-Cts1), and endogenous Cts1 (Koepke et al., 2011). The genetic constructs for Gus-Cts1 and LacZ-Cts1 were stably inserted in the genome at distinct loci resulting in screening strain FB2^CGL^ (Table 1) (Fig. 2A). Strains harboring cytoplasmic Gus or LacZ were used as lysis controls and to mimic defective Cts1 secretion with intracellular reporter accumulation (control strains FB2 Gus_cyt_ and FB2 LacZ_cyt_). Plate assays with the colorimetric substrates X-Gluc (5-bromo-4-chloro-3-indolyl-beta-d-glucuronic acid) and X-Gal (5-bromo-4-chloro-3-indolyl-β-D-galactopyranoside) demonstrated that the screening strain developed the expected blue color indicative for extracellular Gus and LacZ activity while the controls showed no color. Artificial cell lysis demonstrated the presence of intracellular reporter activity in the controls confirming the absence of significant cell lysis (Fig. 2B). Furthermore, quantitative liquid assays with the colorimetric substrate ONPG (o-nitrophenyl-β-D-galactopyranoside) and the fluorogenic substrates MUG and MUC (4-methylumbelliferyl-ß-D-glucuronide trihydrate; 4-methylumbelliferyl β-D-N,N′,N′′-triacetylchitotrioside hydrate) detecting extracellular LacZ, Gus and Cts1 activity, respectively, revealed strongly enhanced extracellular activities of all reporters in FB2^CGL^ compared to the respective control strains containing cytoplasmic versions or lacking the reporters (Fig. 2C). Similar activity measurements in cell extracts again confirmed that all reporters were functional (Fig. 2D). In addition, since we intended to use LacZ activity for high-throughput screening, a mixing experiment to assay its suitability was conducted. Mixing of FB2^CGL^ with FB2 LacZ_cyt_ in different ratios on indicator plates containing X-Gal showed that the ratio of mixing was reflected by the ratio of colonies with a blue halo versus colorless colonies (Fig. 2E-G). This demonstrated that mutants with defective secretion can be identified vis-a-vis colonies with intact secretion as an important requirement for the screening procedure. Finally, plant infection experiments indicated that pathogenicity of FB2^CGL^ was not impaired, since FB2^CGL^ (*a2b2*) crossed with the compatible mating partner FB1 (*a1b1*) elicited the typical symptoms including tumor formation on maize seedlings (Fig. S3). Thus, genetic back-crossing experiments are feasible with this strain. In summary, we successfully designed the strain FB2^CGL^ for screening mutants defective in unconventional secretion of Cts1.

**Fig. 2.**
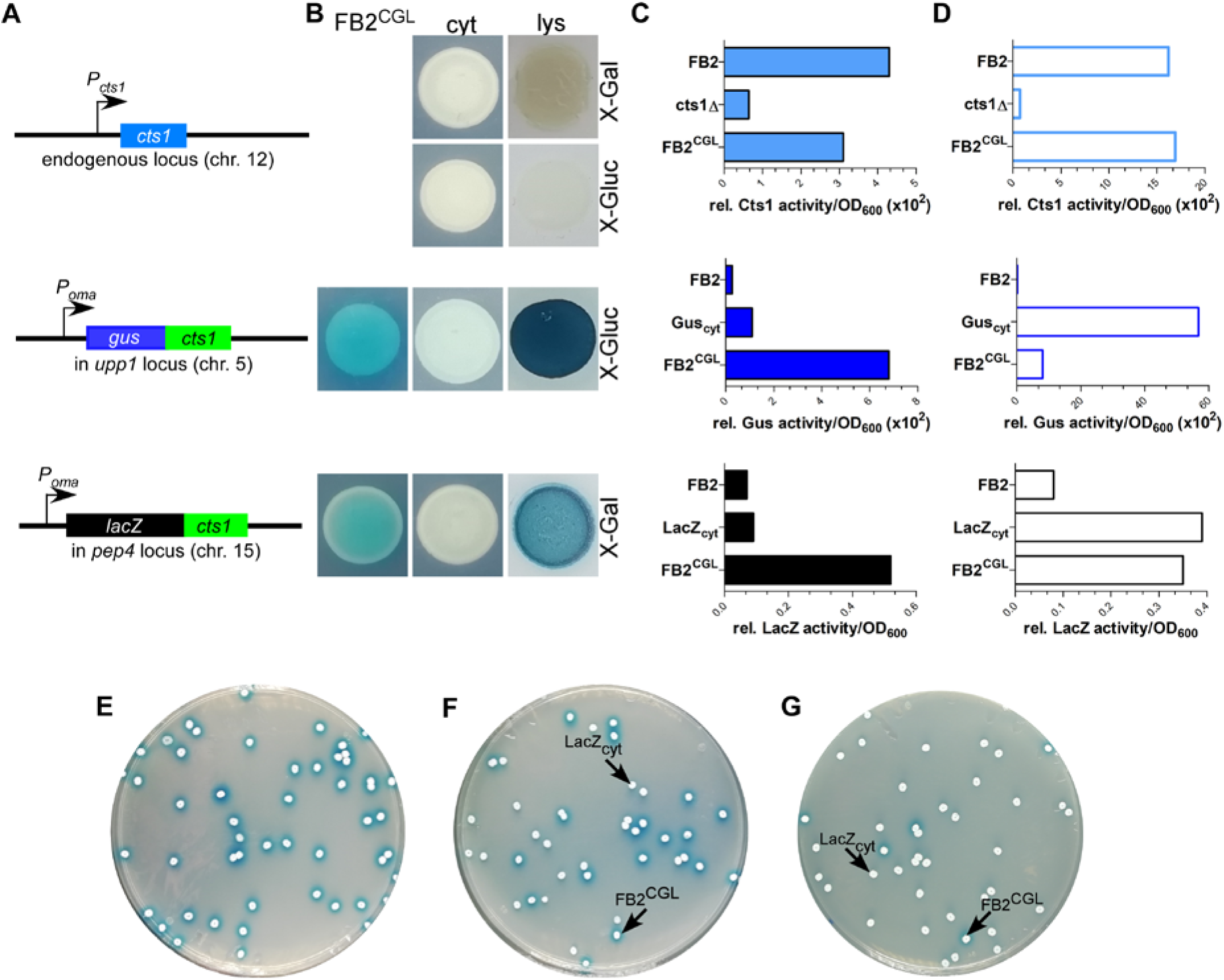
Establishing screening strain FB2^CGL^ harboring three reporters for unconventional secretion. **(A)** Scheme of the genetic constructs for the three reporters present in distinct loci of *U. maydis* strain FB2^CGL^. The reporter genes encoding endogenous Cts1 as well as Gus-Cts1 and LacZ-Cts1 are located on three different chromosomes (chr.). While the *cts1* gene on chr. 12 has not been modified and is present in its natural setup controlled by its native promoter active in yeast cells, both *lacZ:cts1* and *gus:cts1* gene fusions were inserted artificially by homologous recombination using described protease loci (16). Both translational fusion genes are hooked up to the strong synthetic promoter P_*oma*_ which is constitutively active in yeast cells grown on CM-glc. Insertion of the two reporters results in deletion of the genes *upp1* and *pep4* which both encode harmful extracellular proteases (16). **(B)** Plate assay to determine the suitability of the used reporters in FB2^CGL^ in comparison to the lysis controls FB2 Gus_cyt_ and FB2 LacZ_cyt_ harboring cytoplasmic reporter enzymes. FB2 was used as negative control containing neither Gus nor LacZ. X-Gluc and X-Gal were applied as colorimetric substrates for Gus and LacZ, respectively. To visualize intracellular reporter activity, cells were lysed using liquid nitrogen. cyt, control strains with intracellular reporter activity or absent reporter activity. lys, cells lysed by treatment with liquid nitrogen. **(C)** Liquid assays using the substrates MUG, ONPG and MUC to determine the suitability of the three reporters Gus-Cts1, LacZ-Cts1 and Cts1, respectively. 10 μg cell extracts of the indicated strains were used to assay intracellular reporter activities. Upper panel: Cts1 activity based on conversion of MUC; middle panel: Gus activity based on conversion of MUG; lower panel: LacZ activity based on conversion of ONPG. Error bars represent standard deviation of three biological replicates. **(D)** Liquid assays using the substrates MUG, ONPG and MUC to determine the suitability of the three reporters Gus, LacZ and Cts1, respectively. Culture supernatants (Gus-Cts1/LacZ-Cts1) or intact cells (Cts1) of indicated strains were tested to determine extracellular reporter activities. Upper panel: Cts1 activity based on conversion of MUC; middle panel: Gus activity based on conversion of MUG; lower panel: LacZ activity based on conversion of ONPG. Error bars represent standard deviation of three biological replicates. **(E-G)** Mixed culture experiments to verify applicability of the LacZ reporter for high-throughput screening on the colorimetric substrate X-Gal. Screening strain FB2^CGL^ and control strain FB2 LacZ_cyt_ were mixed in the indicated rations and plated onto indicator plates containing X-Gal to visualize extracellular LacZ activity (blue color). Photographs were taken after incubation for 1 d. **(E)**100% FB2^CGL^; **(F)** 50% FB2^CGL^/50% FB2 LacZ_cyt_; **(G)** 10% FB2^CGL^/90% FB2 LacZ_cyt_.

In order to screen for diminished unconventional Cts1 secretion, strain FB2^CGL^ was subjected to UV irradiation with approximately 1% survival rate (Fig. 1, Fig. S2). Mutagenized cells were plated on X-Gal plates to detect extracellular LacZ activity based on the reporter LacZ-Cts1. Approximately 185,000 colonies were screened with a focus on mutants that exhibited normal growth (i.e. normal colony size) but impaired Cts1 secretion. 2,087 candidate mutants showing strongly reduced or absent blue halos were patched on X-Gluc plates employing the Gus marker (Fig. 1). Of those, 566 that stayed colorless again were retested in qualitative plate assays using both the LacZ and the Gus marker to confirm these results. To this end the mutant candidates were patched each on an X-Gluc and an X-Gal containing plate and the coloration was observed. 112 remaining colorless candidates were assayed for Cts1, Gus and LacZ activity in quantitative liquid assays using the substrates MUC, MUG and ONPG, respectively. The different enzyme activities of the progenitor strain FB2^CGL^ were used as a baseline and set to 100%. Again, mutants showing reduced growth in liquid culture were sorted out to ensure that reduced secretion is not connected to growth problems. Multiple mutants displayed slight reduction in the extracellular activity of the reporters. In addition, three mutants were identified, in which Cts1, LacZ and Gus activity was present intracellularly, but diminished extracellularly (below 20% residual activity in all cases; FB2^CGL^mut1-3; Table 1; Fig. 3A,B). Western blot analysis confirmed equal protein amounts for LacZ-Cts1 and Gus-Cts1 in cell extracts of these mutants (Fig. S4A). All mutants showed wildtype growth rates suggesting that reduced secretion is not resulting from growth defects (Fig. S4B).

**Fig. 3.**
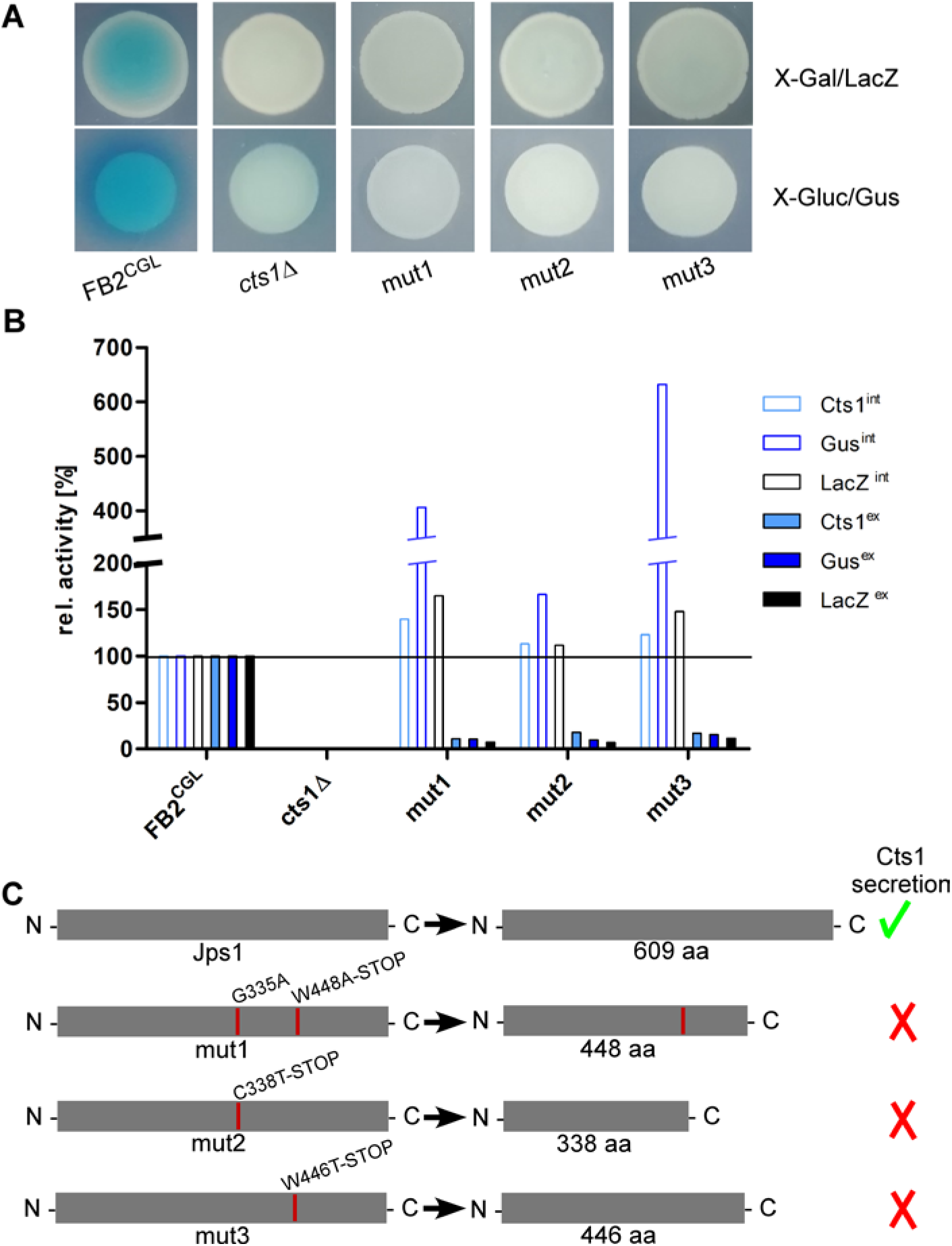
The screen identifies the uncharacterized protein Jps1. **(A)** Plate assays of indicated strains for extracellular LacZ activity using X-Gal and extracellular Gus activity using X-Gluc. The assays are based on secretion of the reporters LacZ-Cts1 and Gus-Cts1 and the respective substrates are only converted, if the fusion protein is secreted. This leads to the formation of blue colonies (and blue halos in case of Gus) resulting from extracellular substrate conversion. Mutants FB2CGLmut1, mut2 and mut3 were identified after UV mutagenesis of strain FB2CGL on screening plates (white colonies on both X-Gluc and X-Gal) and show the expected reduced extracellular activities indicating deficiency in Cts1 secretion. **(B)** Quantitative liquid assays detecting intracellular (int, filled columns) and extracellular (ex, open columns) Gus, LacZ and Cts1 reporter activity. Screening strain FB2CGL and a *cts1* deletion mutant (AB33 cts1Δ, 15) were used as positive and negative controls, respectively. Activities obtained for FB2CGL were set to 100% to allow for a direct comparison of all strains. The assay was conducted thrice with similar results and a representative replicate is shown. **(C)** In all three identified mutants, base exchanges in gene *umag_03776*, now termed *jps1*, were identified. At the aa level, mutations result in the introduction of premature stop codons leading to the production of truncated proteins in all three mutants. Mut1 carries an additional aa exchange at position 335. Mutations identified in FB2CGLmut1-3 are indicated with red lines in the schematic representation of the protein. The size of native Jps1 is 609 aa.

To identify the responsible mutations whole genome sequencing comparing FB2^CGL^mut1 with its progenitor FB2^CGL^ was performed, using the published sequence of *U. maydis* UM521 as a template for the assembly (https://mycocosm.jgi.doe.gov/Ustma2_2/Ustma2_2.home.html; last accessed 2020/04/01). 32 base-pair substitutions were detected in the comparison of FB2^CGL^ and FB2^CGL^mut1. 9 single nucleotide polymorphisms (SNPs) were located in non-coding regions and were thus unlikely to cause the observed defect in unconventional secretion. The majority of the 23 mutations in coding regions were found to be 5’-C→T or 5’-CC→TT transitions which are expected for UV-induced mutations (Fig. S5) (Pfeifer et al., 2005). 11 substitutions were irrelevant silent mutations. The remaining 12 SNPs led to 10 aa replacements in 9 encoded proteins and hence constituted the top remaining candidates (Fig. S5). To locate which of these mutations was responsible for the defective Cts1 secretion, genetic back-crossing experiments were performed using plant infections. FB2^CGL^mut1 was pathogenic in crosses with compatible FB1^CGL^ which carries similar reporter genes but an opposite mating type (Fig. S1; Fig. S3; see Materials and Methods). Meiotic progeny was obtained and assayed for extracellular Gus, LacZ and Cts1 activity on indicator plates and by quantitative liquid assays. Based on these results the progeny was grouped into mutants with defective and intact secretion (Fig. S6). Sequencing of the 9 candidate genes obtained from comparative genome sequencing revealed that all progeny with reduced extracellular reporter activity harbored mutations in gene *umag_03776* (Fig. 3, Fig. S6). Strikingly, also strains FB2^CGL^mut2 and mut3 carried detrimental mutations in *umag_03776*, leading to synthesis of C-terminally truncated proteins (Fig. 3C). This strongly suggested that this gene is essential for Cts1 secretion. The corresponding gene product was subsequently termed Jps1 (jammed in protein secretion screen 1). While the protein Jps1 has a predicted length of 609 aa, the truncated versions produced in mutants FB2^CGL^mut1, mut2 and mut3 only contained 448, 338 and 446 aa, respectively (Fig. 3C). In essence we identified a crucial factor for Cts1 secretion by genetic screening.

### Jps1 is essential for Cts1 localization and secretion

Jps1 is annotated as hypothetical protein with unknown function (https://mycocosm.jgi.doe.gov/Ustma2_2/Ustma2_2.home.html; last accessed 2020/04/01) and does not contain yet known domains (SMART; http://smart.embl-heidelberg.de/; last accessed 2020/04/01). None of its homologs in other Basidiomycetes has been characterized so far and Ascomycetes like *S. cerevisiae* lack proteins with significant similarities. Thus, to obtain first insights into its function we initially validated the screen by deleting the respective gene in the background of laboratory strain AB33 (Brachmann et al., 2001) using homologous recombination (AB33jps1Δ). Microscopic analysis revealed that budding cells of the deletion strain had a normal morphology comparable to controls like AB33 or the chitinase deletion strain AB33cts1Δ (Fig. 4A) (Langner et al., 2015). Chitinase assays showed strongly reduced extracellular activity, confirming the essential role of Jps1 for Cts1 secretion (Fig. 4B). To analyze the localization of Cts1 in a *jpsΔ* background strain AB33 jps1Δ/CtsG expressing a functional Cts1-Gfp (Cts1G) fusion was generated (Fig. 4C) (Koepke et al., 2011). Microscopic studies revealed that in contrast to its native localization in the fragmentation zone (Fig. 4D) (Langner et al., 2015; Aschenbroich et al., 2019) the Gfp signal for Cts1G now accumulated intracellularly and at the septa (Fig. 4E,F). Interestingly, in about 56% of the cases, the signal was detected at the primary septum only, while in the remaining 44% the signal was present at both septa (Fig. 4G). Cts1G was never observed in the fragmentation zone of the *jps1* deletion strain (Fig. 4H).

**Fig. 4.**
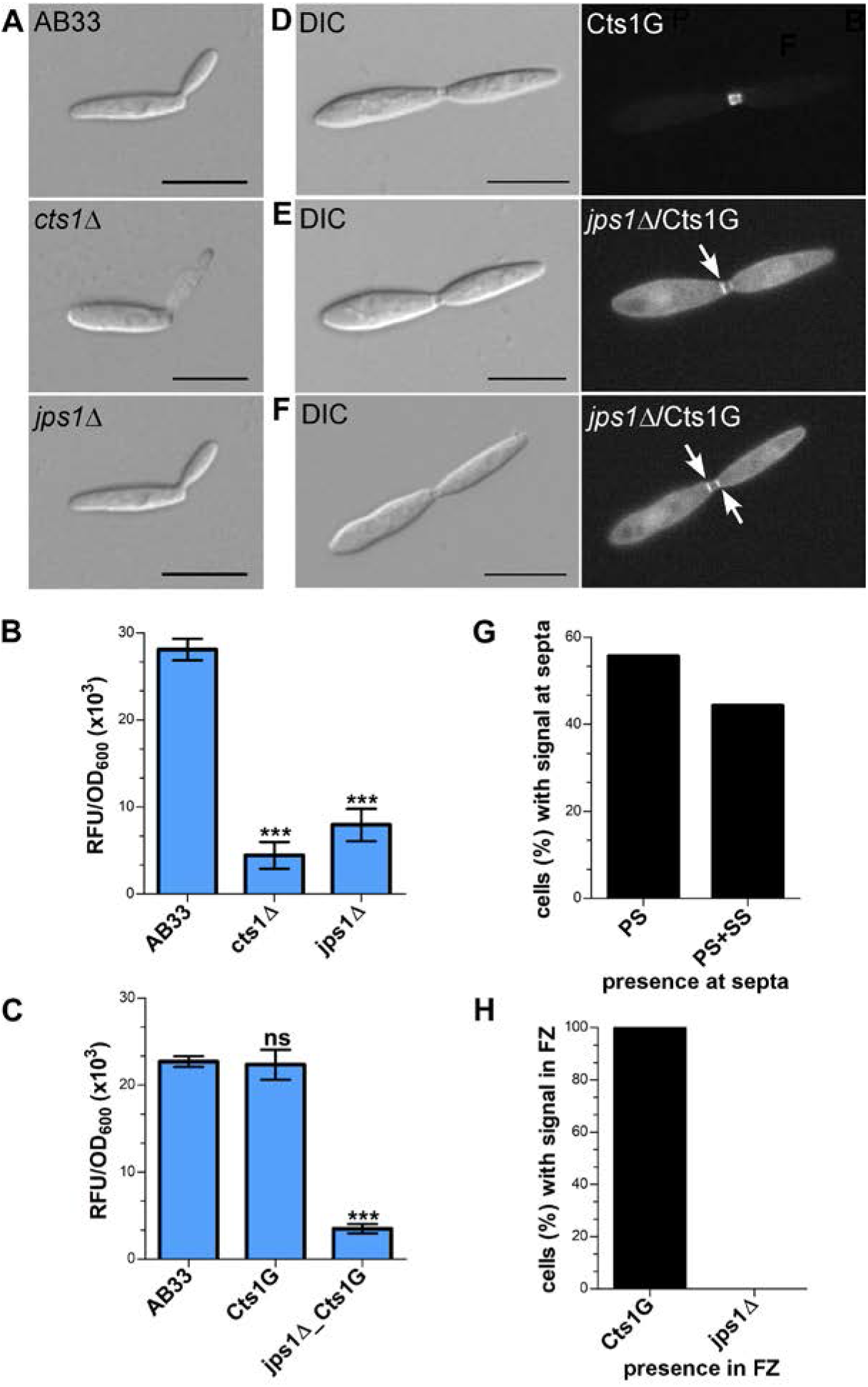
Jps1 is crucial for unconventional Cts1 secretion. **(A)** Micrographs of yeast-like growing cells of indicated strains. The *jps1* deletion strain AB33jps1Δ (jps1Δ) does not show any morphological abnormalities. The *cts1* deletion strain AB33cts1Δ (cts1Δ) and the progenitor laboratory strain AB33 (wt) are shown for comparison. Scale bars, 10 μm. **(B)***jps1* deletion in the AB33 verifies its essential function in Cts1 secretion. Extracellular Cts1 activity of AB33, AB33jps1Δ and the control AB33cts1Δ is depicted. The assay was conducted in three biological replicates. Error bars indicate standard deviation. ***, p value 0.001; n.s., not significant (two sample t-test). **(C)** Extracellular Cts1 activity of Cts1G expressing strains are comparable to the progenitor strain AB33, suggesting that the protein is functional. The assay was conducted in five biological replicates. Error bars indicate standard deviation. ***, p value 0.001; n.s., not significant (two sample t-test). **(D-F)** Localization of Cts1-Gfp (Cts1G) in AB33 **(D)** and AB33jps1Δ **(D,E)**. While Cts1 accumulates in the fragmentation zone of dividing cells with two septa in AB33 **(D)**, it enriches in the cytoplasm and at the septa in the *jps1* deletion strain. Two different scenarios were observed: Either Cts1 was found only at the primary septum at the mother cell side **(E)** or at both septa **(F)**. White arrows depict septa with Cts1 signal. Scale bars, 10 μm. **(G)** Distribution of Cts1G signal at the primary septum only (PS) and at both septa (PS+SS) of dividing AB33jps1Δ cells with completely assembled fragmentation zones (AB33Cts1G: 900 cells analyzed; AB33jps1Δ/Cts1G: 1370 cells analyzed; three biological replicates). **(H)** Cts1G is restricted from the fragmentation zone. The graph depicts the fraction of cells in exponentially growing cultures of indicated strains with Cts1G accumulation in the fragmentation zone (similar cells analyzed as shown in G).

Next, a strain expressing Jps1 fused to Gfp (Jps1G) was generated to localize the protein (AB33Jps1G). Chitinase assays verified the functionality of the fusion protein (Fig. 5A). Intriguingly, microscopic analysis revealed that Jps1G accumulated in the fragmentation zone of budding cells, similar to Cts1 (Fig. 5B). To investigate if Jps1 localization depends on chitinase function, Cts1 was deleted in the background of AB33JpsG (AB33cts1Δ/Jps1G). Interestingly, this did not disturb Jps1 localization. This suggests that Cts1 is dispensable for Jps1 function but not *vice versa*, indicating a unidirectional dependency of Cts1 on Jps1. To further substantiate the apparent co-localization, both proteins were differentially tagged in a single strain, expressing Jps1 fused to mCherry and Cts1 fused to Gfp (AB33Cts1G/Jps1mC). Indeed, both signals completely overlapped in the fragmentation zone in each observed case (Fig. 5D, E). Since the co-localization studies suggested that the two proteins might interact we conducted yeast two-hybrid assays in which we fused Jps1 with the activation domain (AD) and Cts1 with the binding domain (BD), and *vice versa*. Self-interaction could be detected neither for Jps1 nor for Cts1. However, for one of the combinations, a weak interaction between Jps1 and Cts1 could be observed in serial dilutions on selection plates, indicating that the two proteins might interact (Fig. 5F).

**Fig. 5.**
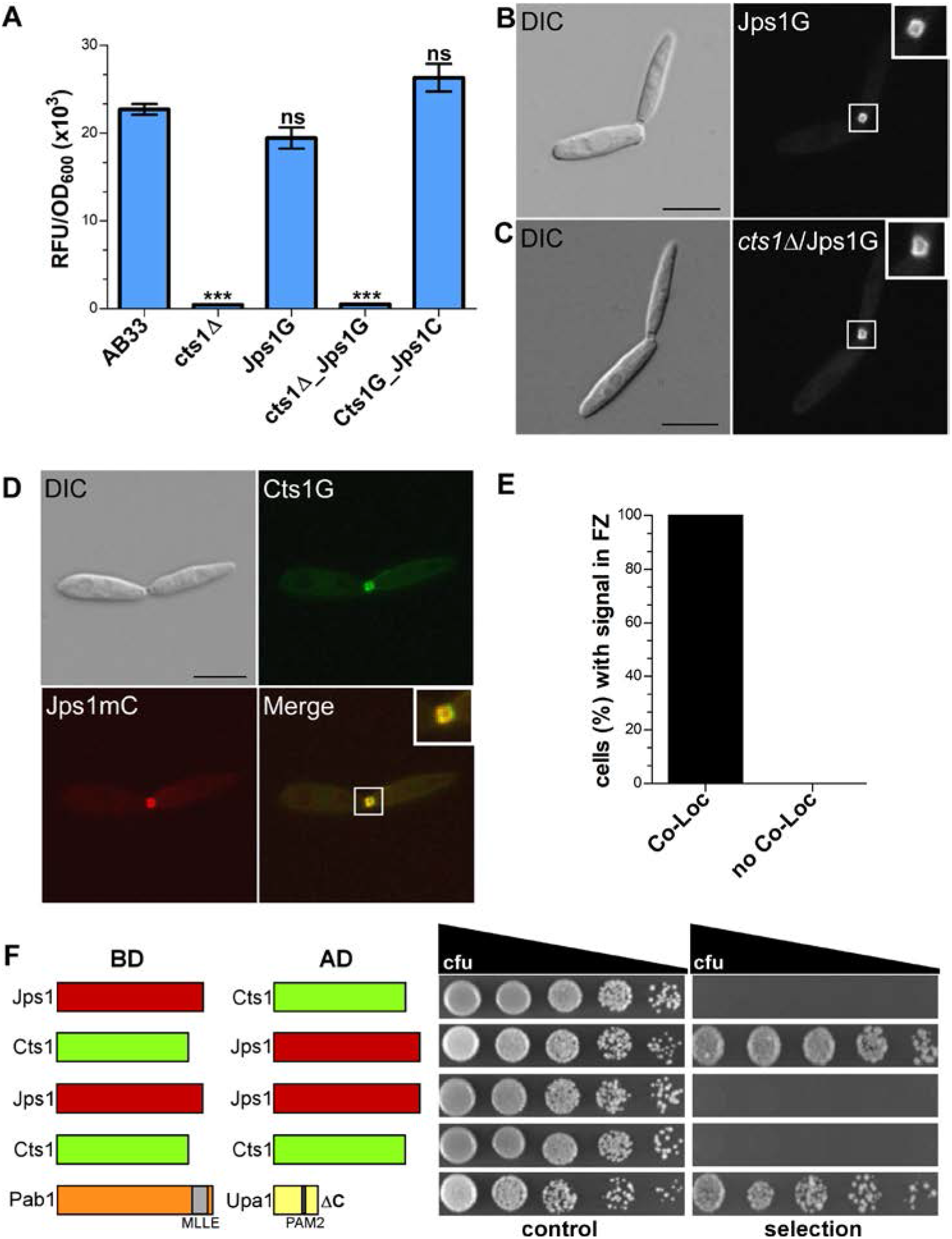
Jps1 co-localizes with Cts1 in the fragmentation zone. **(A)** Extracellular Cts1 activity of indicated strains. AB33cts1Δ lacking Cts1 was used as negative control. The Jps1mC fusion protein is functional. The assay was conducted in five biological replicates. Error bars indicate standard deviation. ***, p value 0.001; n.s., not significant (two sample t-test). **(B)** Localization of Jps1G in AB33. The protein accumulates in the fragmentation zone of dividing cells. Scale bar, 10 μm. **(C)** Localization of Jps1G in AB33cts1Δ. Localization of Jps1 in the fragmentation zone is not altered in the absence of Cts1. Scale bar, 10 μm. **(D)** Micrographs of strain AB33Cts1G/Jps1mC indicating co-localization of Cts1G and Jps1mC in the fragmentation zone. **(E)** Quantification of co-localizing signals (Co-Loc) and not co-localizing signals (No Co-Loc) of strain AB33Cts1G/Jps1mC in the fragmentation zones of 970 cells observed. The experiment was conducted in 3 biological replicates. **(F)** Yeast-two hybrid assays to analyze protein:protein interactions between Jps1 and Cts1. For the positive control using strains producing Pab1 and Upa1 a weak interaction had been shown before (Pohlmann et al., 2015). BD, binding domain; AD, activation domain.

Thus, our genetic screen identified a novel essential protein for unconventional Cts1 secretion. Co-localization and interaction between Jps1 and Cts1 suggest that Jps1 might act as an anchoring factor for Cts1 that supports its local accumulation in the fragmentation zone.

## 4. Discussion

In this study we identified Jps1, a novel factor essential for unconventional export of chitinase Cts1 via the fragmentation zone of budding cells. Jps1 was identified in a forward genetic screen. Such genetic screens are powerful tools to identify important players in unknown pathways (Forsburg, 2001). The prime example is the screen for components of the conventional secretion pathway which has been performed in *S. cerevisiae*. Initially, this temperature-sensitive screen was based on the fact that proteins accumulate in the endomembrane system of cells in which secretion is disturbed. These dense cells can be separated from cells with intact secretion by gradient centrifugation (Novick et al., 1980). The screen was continuously further developed and finally provided a detailed view on the key components of the canonical pathway including all stages of protein export (Mellman and Emr, 2013). Similarly, the here employed screen proved to be very efficient and powerful: in all three obtained mutants, mutations localized to the same gene (*jps1*). On the one hand, this underlines the quality of the screening procedure. On the other hand, this observation may also limit the screen in that other factors may be hard to identify in this screening set up. Alternatively, the repeated identification of the same mutant could be due to the fact that there is only one major factor involved in Cts1 secretion. However, we consider that unlikely. Therefore, in the next step a second copy of Jps1 under its native promoter will be inserted into the genome. This will minimize the risk of identifying the gene again in further screening attempts. Furthermore, we will streamline identification of responsible mutations. Here, we used genetic back-crosses via plant infection in combination with a PCR approach to identify Jps1. For future studies, we will use batch-sequencing of mutants with or without secretion of the reporters (Fig. S2 steps 10 & 11). Such pooled linkage analysis based on next-generation sequencing is known to have a great statistical power and thus allows an efficient identification of underlying mutations (Birkeland et al., 2010).

In the first screening round we concentrated on mutants that showed a normal growth behavior (i.e. colony sizes similar to untreated cells after UV mutagenesis). Mutants that are impaired in growth are much more complicated to analyze. Discrimination between true secretory defects and reduced secretion due to poor fitness of the cells is very difficult. Hence, it is well conceivable that we missed other important factors, especially because unconventional Cts1 secretion is tightly connected to cytokinesis (Aschenbroich et al., 2019). This also explains why we did not identify mutants defective in the septation factors Don1 or Don3 which we have shown to be essential for unconventional Cts1 secretion (Aschenbroich et al., 2019). The cognate deletion mutants have a cytokinesis defect and grow in tree-like structures (Weinzierl et al., 2002).

As an alternative application, the screen will be employed to optimize our recently established protein expression platform (Feldbrügge et al., 2013; Sarkari et al., 2016). Here we use Cts1 as a carrier for valuable heterologous proteins. Exploiting the unconventional secretion route brings the advantage that *N-*glycosylation is circumvented and thus, sensitive proteins like bacterial enzymes can be exported in an active state (Sarkari et al., 2014; Terfrüchte et al., 2017; Stoffels et al., 2019). The screen will be adapted to select for mutants with enhanced marker secretion to eliminate existing bottlenecks and thus enhance yields (Terfrüchte et al., 2018). Random mutagenesis screens for hypersecretors were for example key to establish industrial production strains like the cellulase producing filamentous fungus *Trichoderma reesei* strain RutC-30 in which amongst further changes carbon catabolite repression was eliminated (Peterson and Nevalainen, 2012). A restriction enzyme mediated insertion (REMI) screen based on a β-galactosidase reporter has also led to the discovery of a set of mutants showing supersecretion of the reporter in *Pichia pastoris* (Larsen et al., 2013).

Our findings confirm the proposed lock-type secretion mechanism (Aschenbroich et al., 2019; Reindl et al., 2019) and indicate that we indeed discovered a novel pathway of unconventional secretion. Jps1 supports the subcellular accumulation of Cts1 in the fragmentation zone from where it is released. While the secretory pathway itself is new, it is conceivable that the molecular details of Cts1 export might be similar to described systems. For example, it could be released via self-sustained translocation through the plasma membrane which has been described for FGF2, but is nowadays discussed also for other proteins like HIV-Tat and interleukin 1beta (Dimou and Nickel, 2018). In the future, biochemical studies will shed light on the molecular pathway of unconventional Cts1 secretion. Interestingly, Jps1 orthologs are restricted to the Basidiomycetes. Hence, lock-type secretion is likely conserved at least in the Ustilaginales like *Sporisorium reilianum*, *Ustilago hordei*, *Pseudozyma aphidis* or *Tilletia walkeri*. This assumption is supported by the finding that Cts1 orthologs lacking predictions for N-terminal signal peptides are also present in these species. Unfortunately, published information about the yeast-like growth of these fungi and potential formation of fragmentation zones is yet scarce. The absence of Jps1 orthologs in *S. cerevisiae* and other Ascomycetes, fits well to the observation that septation in *S. cerevisiae* does not involve formation of a fragmentation zone, suggesting that molecular details of cell division differ between the two organisms (Reindl et al., 2019).

Jps1 is essential for efficient Cts1 localization and secretion while in turn Cts1 is not needed for Jps1 accumulation in the fragmentation zone. This one-sided dependency indicates that Jps1 is an important factor for Cts1 secretion but not *vice versa*. The underlying molecular details and thus the exact role of Jps1 during Cts1 secretion remain to be addressed by detailed biochemical and cell biological studies in the future. It is also not clear how the two proteins reach the fragmentation zone. Moving early endosomes enrich in the fragmentation zone prior to budding and were shown to carry Don1 (Weinzierl et al., 2002; Schink and Bölker, 2009). They are thus prime candidates for transporting proteins into the small compartment. However, we neither observed Cts1 nor Jps1 on these motile organelles (data not shown). While the septation factors Don1 and Don3 seem to play a passive role for Cts1 release by sealing off the fragmentation zone with the secondary septum, Jps1 acts as a third factor for lock-type unconventional secretion which is likely directly involved in the export process. Based on our results, it is conceivable that Jps1 functions as an anchoring factor for Cts1, thus supporting its local accumulation in the fragmentation zone where it likely degrades remnant chitin together with conventionally secreted Cts2 to support cytokinesis (Fig. 6).

**Fig. 6.**
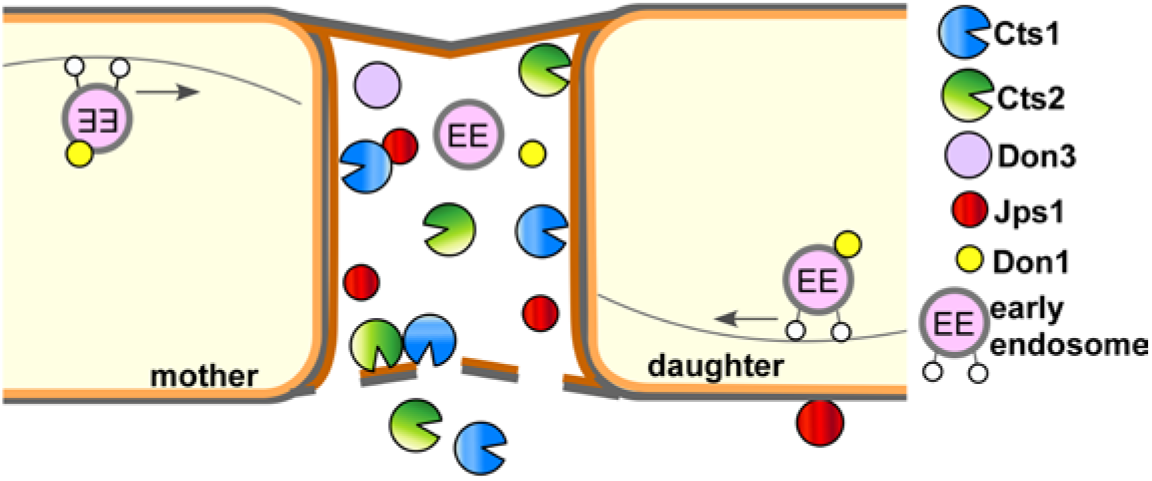
Current model for subcellular targeting and unconventional secretion of Cts1 via anchoring factor Jps1. Cts1 is targeted to the fragmentation zone via a potential lock-type unconventional secretion mechanism. Motile early endosomes shuttle bidirectionally through the cells and transport the septation factor Don1 which is essential for secondary septum formation. Together with Don3 it localizes to the fragmentation zone formed between mother and daughter cell during cytokinesis. Both Don1 and Don3 are crucial for Cts1 export. The newly identified factor Jps1 also accumulates in the fragmentation zone. We hypothesize that the protein functions in anchoring Cts1 in the small compartment. Here, Cts1 acts in degrading remnant chitin for detaching mother and daughter cell in concert with conventionally secreted Cts2.

In sum, our genetic screen has already proven to be very efficient and an improved version will deal as a basis to identify further key components of unconventional secretion and optimize the connected protein expression platform in the future.

## Author Contributions

J.S. designed, conducted and evaluated the genetic screen with support of M.R., K.H. and A.G., M.R. characterized Jps1 and prepared micrographs, quantifications and enzyme assays. A.B. performed genome sequencing. K.S. prepared the manuscript with input of all co-authors, directed the project and acquired funding.

## Acknowledgments

We acknowledge Dr. M. Feldbrügge for continuous support and valuable discussion. We thank B. Axler and U. Meyer for excellent technical support of the project. T.E. Hyland and L. Mielke contributed to the project in the framework of an internship and M. Tulinski and S. Wolf in their Master projects.

