## Supplementary Material for "A novel factor essential for unconventional secretion of chitinase Cts1"

The Supplementary Material contains Supplementary Figures 1-7.

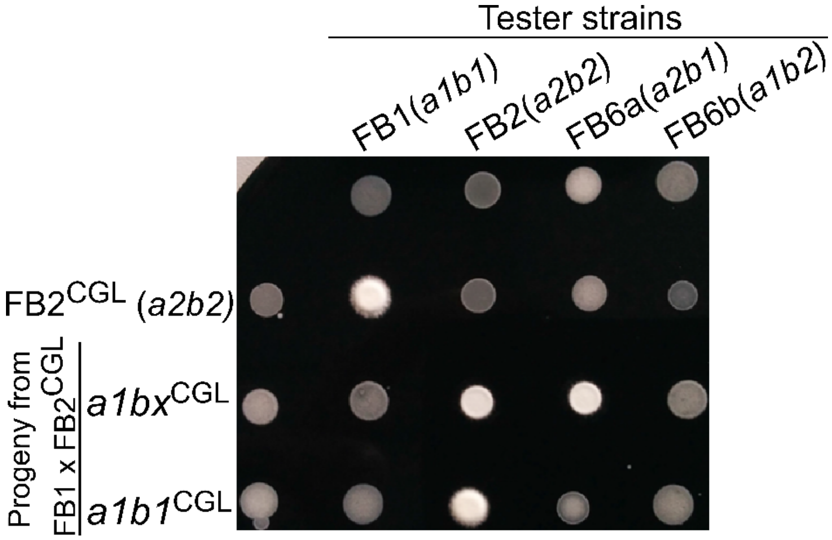


**Supplementary Figure 1.** Mating assay to identify FB1$^{CGL}$ with *a1b1* mating type harboring the reporter LacZ-Cts1 and Gus-Cts1 using charcoal plates. Mating of compatible strains was observed on CM-charcoal plates. Fuzzy colonies indicate mating while smooth colonies show incompatibility. FB1 (mating type *a1b1*), FB2 (mating type *a2b2*), FB6a (mating type *a2b1*) and FB6b (mating type *a1b2*) are tester strains with known genotypes. Progeny carrying the two reporters Gus-Cts1, LacZ-Cts1 was obtained by crossing the screening strain FB2$^{CGL}$ with wild type strain FB1 and tested for mating to identify an *a1b1* clone. One clone of the two displayed clones has an *a1b1* mating type because it shows a fuzzy colony with FB2 but not with FB6a (strain termed FB1$^{CGL}$). This strain was used for genetic back-crosses with mutagenized FB2$^{CGL}$ during the screen.

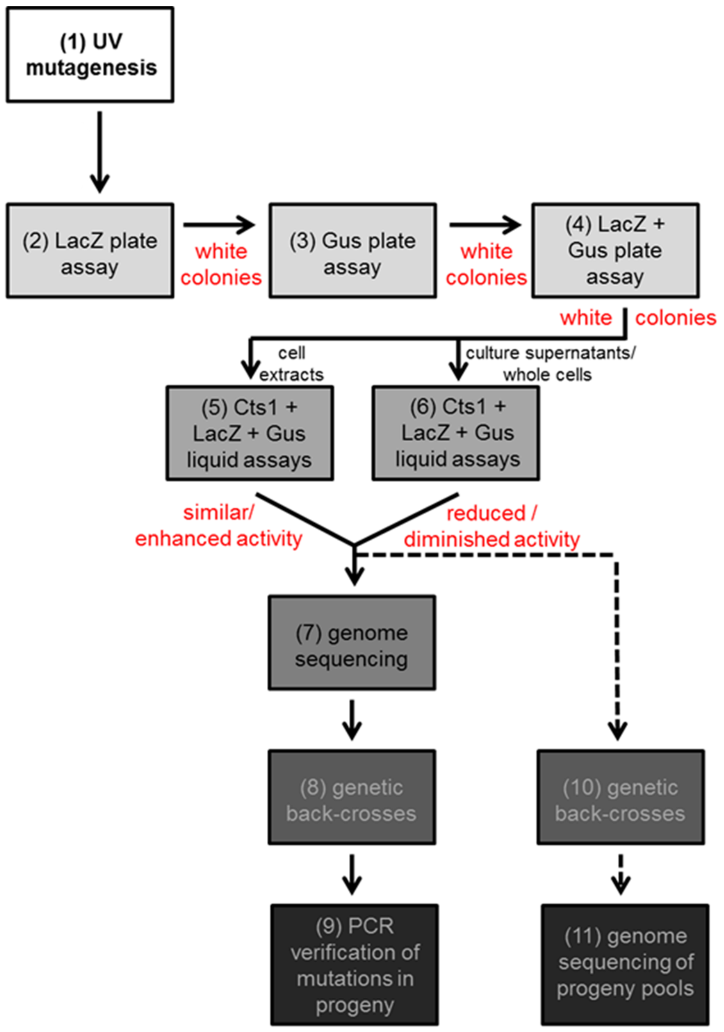


**Supplementary Figure 2.** Detailed step-by-step description of the forward genetic screen. All steps performed to identify factors for unconventional secretion of chitinase Cts1 are indicated in detail. First of all, cells were mutagenized by UV light (1). Next, plate assays were performed (2-4). Initial screening of the mutagenized cells was conducted on X-Gal containing plates (2). White colonies lacking extracellular LacZ activity were then subjected to further plate assays on both X-Gal and X-Gluc (3, 4). Colonies that stayed white on both substrates were analyzed in detail using quantitative liquid assays with substrates for all three unconventional secretion reporters (Cts1, LacZ-Cts1, Gus-Cts1) (5,6). Both intra- (5) and extracellular activities (6) were determined. Genome sequencing and alignment with the progenitor strain FB2$^{CGL}$ revealed multiple mutations (candidate loci) likely resulting from UV treatment (7). Genetic back-crosses with a non-mutagenized FB1 derivative harboring all three reporters (FB1$^{CGL}$) were performed and led to progeny showing defective unconventional secretion while harboring cytoplasmic activity for all three reporters (8). Amplification and sequencing of the candidate loci identified in the alignment identify the responsible mutation (9). In future studies, progeny of the back-crosses could also be pooled according to the absence or presence of unconventional secretion. Genome sequencing of the two pools should also reveal the responsible mutation (10,11).

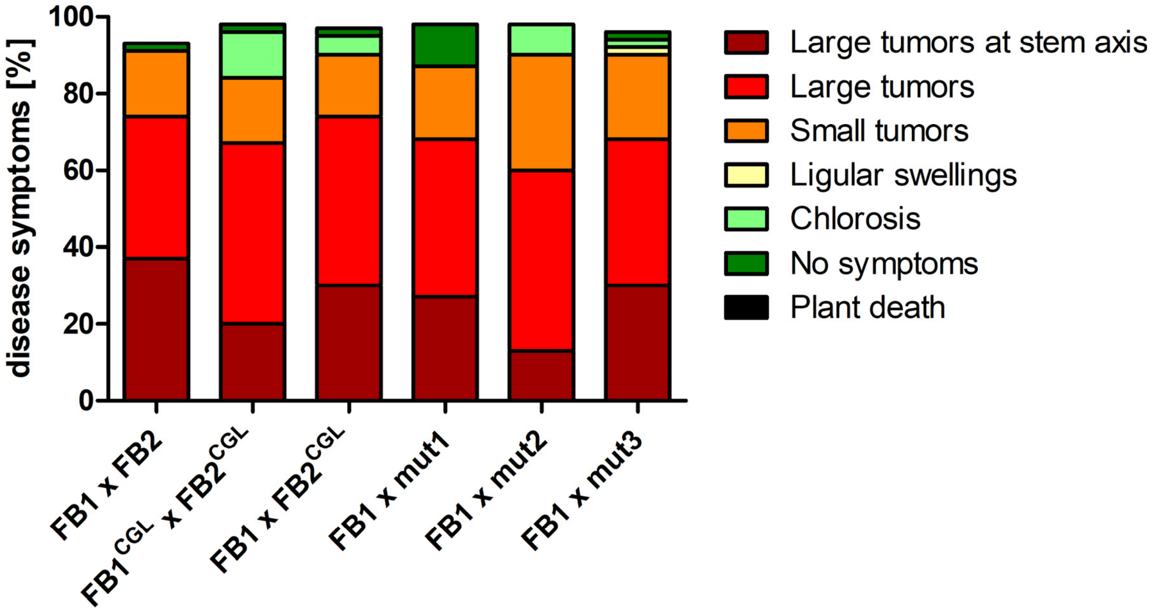


**Supplementary Figure 3.** Plant infections with compatible FB1 and FB2 derivatives. Maize seedlings were inoculated with the indicated compatible strains and disease symptoms scored 7 days post infection. The infection was performed once with 40 plants each. All tested strain combinations were pathogenic and showed symptoms comparable to the wild type cross FB1 (*a1b1*) x FB2 (*a2b2*).

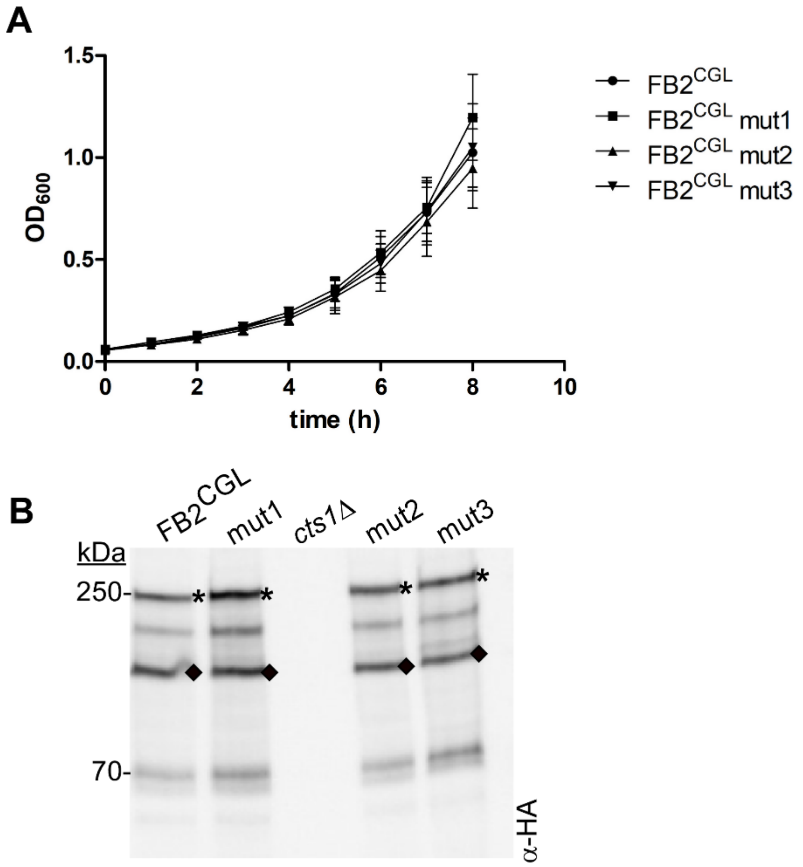


**Supplementary Figure 4.** Mutants impaired in Cts1 secretion grow normal and produce similar levels of Gus-Cts1 and LacZ-Cts1 as the screening strain FB2$^{CGL}$. **(A)** Growth curves of FB2$^{CGL}$ and the indicated UV mutants. Growth was followed by determining the optical density at 600 nm for eight hours. All strains duplicate with comparable rates. **(B)** Western blot detecting Gus-Cts1 and LacZ-Cts1 levels in FB2$^{CGL}$ and the indicated UV mutants. Antibodies directed against the HA-tag in the SHH linker of the fusion proteins were used for detection. Asterisks and rhombs depict the expected sizes of the fusion proteins LacZ-Cts1 and Gus-Cts1, respectively. A strain lacking the reporters was used as negative control (AB33 *cts1*Δ).

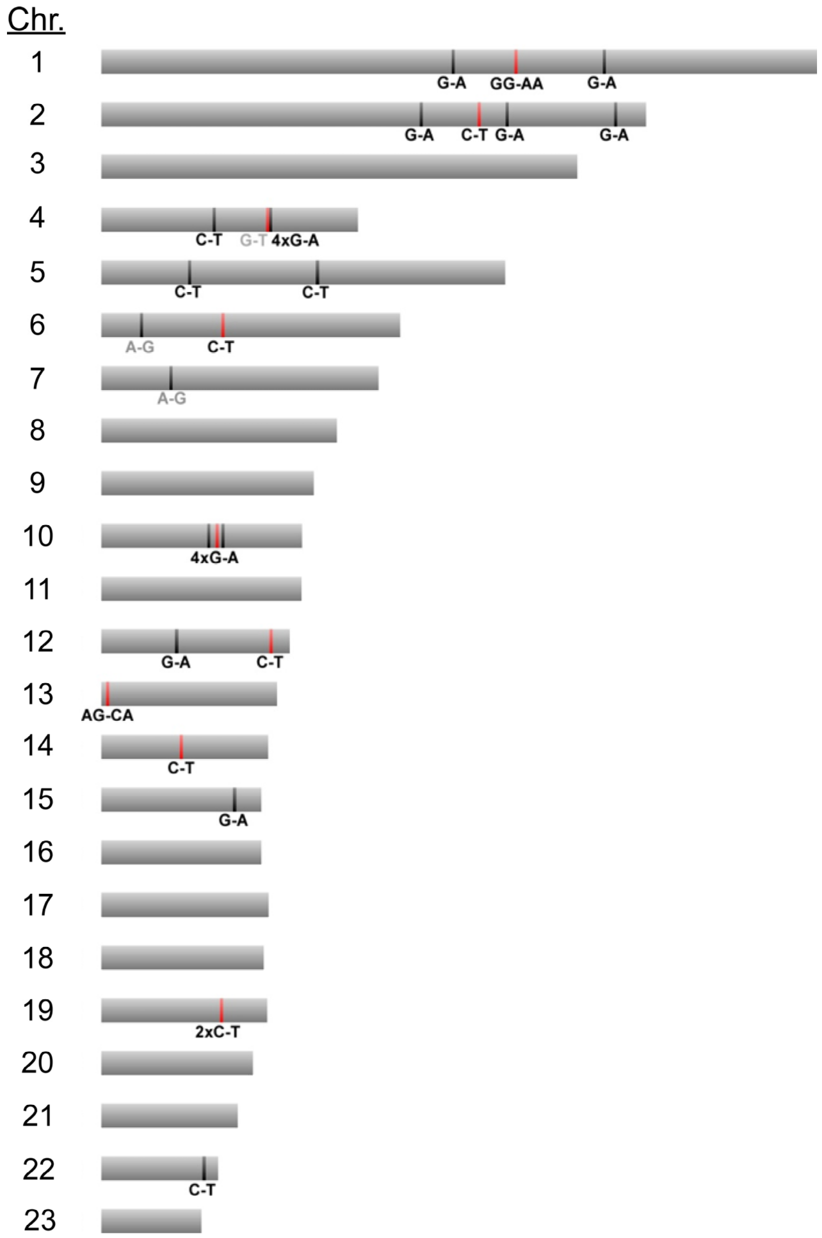


**Supplementary Figure 5.** SNPs identified in genome sequence comparisons between screening strain FB2$^{CGL}$ and mutant candidate FB2$^{CGL}$mut1. The schematic representation shows all 23 chromosomes. Bars depict 32 base changes detected in FB2$^{CGL}$mut1 compared to its progenitor FB2$^{CGL}$. Red bars indicate 12 mutations which go along with amino acid replacements in 9 ORFs and are thus candidates likely responsible for the observed defective secretion. Unexpected base transitions are indicated in grey letters while black letters depict transitions typically observed after UV treatment. Figure dimensions are not to scale.

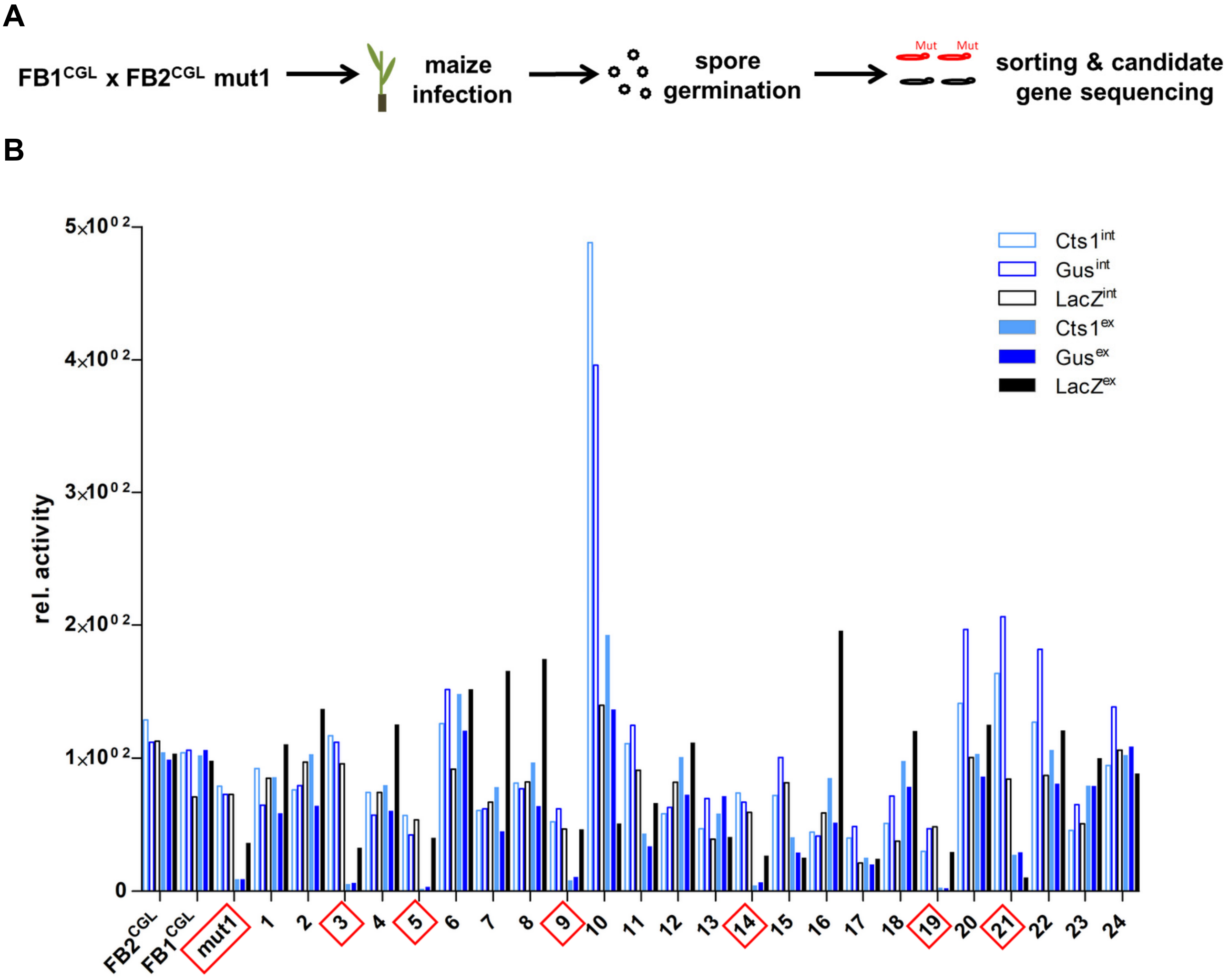


C

| | **Progeny FB1^CGL x FB2^CGL (# artificially numbered)** |
|---|---|
| **Progeny with intact Cts1 secretion** | #1, #2, #4, #6, #7, #8, #10, #11, #12, #13, #15, #16, #17, #18, #20, #22, #24 |
| **Progeny with defective Cts1 secretion** | #3, #5, #9, #14, #19, #21 |

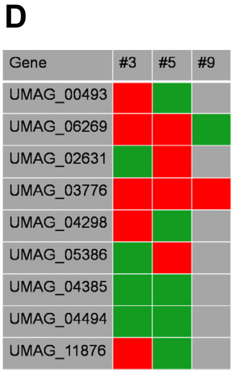


**Supplementary Figure 6.** Extracellular Cts1 activity of progeny from the genetic cross of FB1$^{CGL}$ and FB2$^{CGL}$mut1. **(A)** Back-crossing strategy to allow for identification of responsible mutations in strain FB2$^{CGL}$mut1. Spores obtained after crossing FB1$^{CGL}$ and FB2$^{CGL}$mut1 were germinated and singled out. Single colonies were sorted into two groups (extracellular reporter activity: yes/no) by assaying extracellular reporter activities on X-Gal and X-Gluc plates and by liquid Cts1 assays with whole cells. **(B)** Liquid Gus, LacZ and Cts1 assays to sort progeny obtained from FB1$^{CGL}$ and FB2$^{CGL}$mut1 crosses. Both extracellular (ex, filled columns) and intracellular (int, open columns) activities were determined. Numbers (#) indicate the different clones obtained from germinated spores. The progenitor strains $FB2^{CGL}$, $FB2^{CGL}$mut1 and $FB1^{CGL}$ were used as references for normal and strongly diminished unconventional secretion. Clones with strongly reduced extracellular activities for all three reporters are listed in C and were used for sequencing. **(C)** Progeny groups obtained after genetic back-crossing based on the results shown in **(B)**. **(D)** Sequencing of diagnostic PCR products was conducted on progeny which did not show extracellular reporter activity using primers that encompass the 12 potential mutations leading to aa exchanges in 9 ORFs. Sequencing revealed mutagenized *umag_03776* to be responsible for the reduction in unconventional secretion. Red: candidate locus mutagenized; green: candidate locus wildtypic.

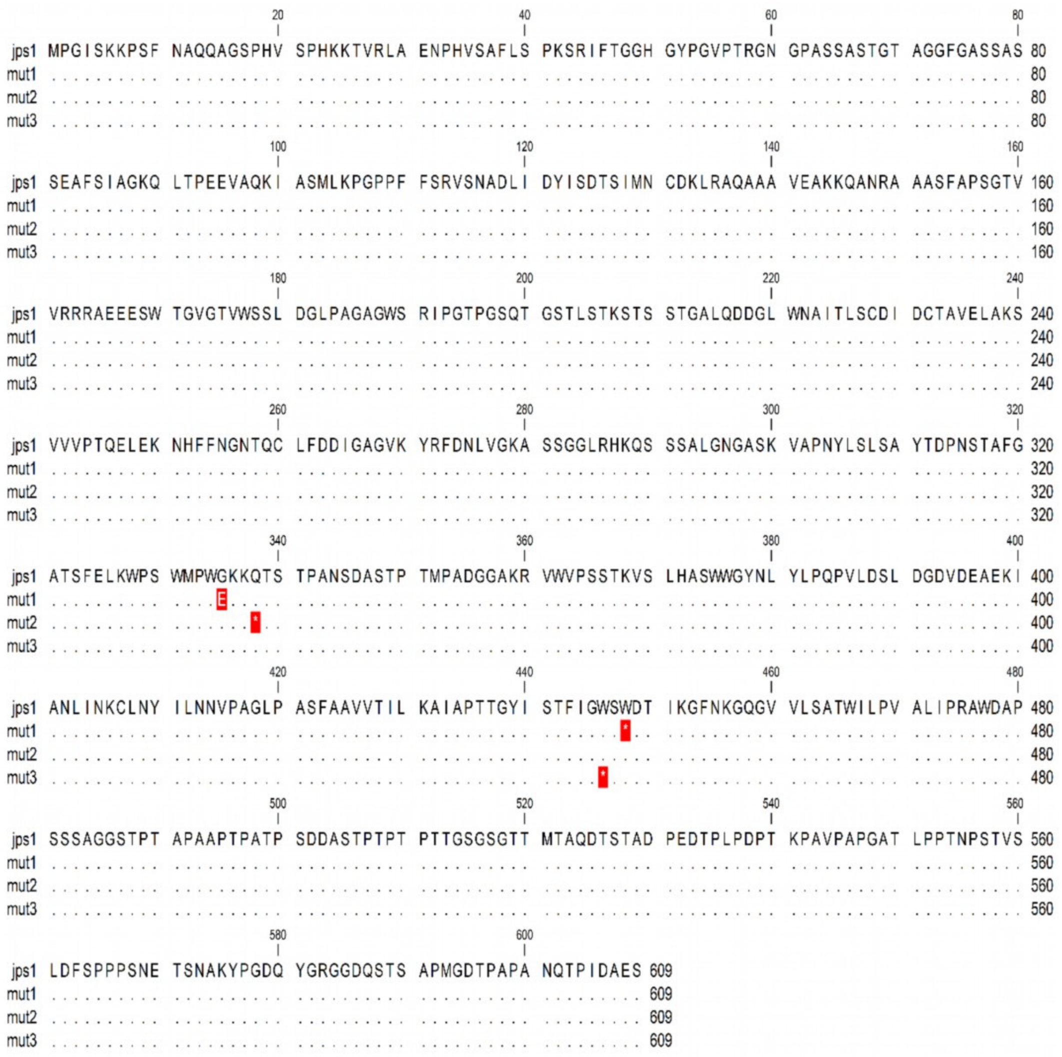

**Supplementary Figure 7.** Aa alignment of Jps1 wild type protein and three mutant versions obtained in the UV mutagenesis for candidates with diminished Cts1 secretion. Red background color indicates aa changes in the mutants in comparison to the native protein (upper line). Asterisks depict stop codons. Mutations result in the production of C-terminally truncated proteins (compare Fig. 3). Numbers show base pair counts.
